# Granularity of thalamic head direction cells

**DOI:** 10.1101/2025.09.08.674912

**Authors:** Sara Hijazi, Shan Jiang, Mara S. Wülfing, Jacqueline Quach, Patrick A. Lachance, Michael E. Hasselmo, Tim J. Viney

## Abstract

Head direction signaling is fundamental for spatial orientation and navigation. The anterodorsal nucleus of the thalamus (ADn) contains a high density of head direction (HD) cells that process sensorimotor inputs for subsequent synaptic integration in postsynaptic cortical areas. We tested the hypothesis that individual HD cells show differences in their firing patterns and connectivity by recording and juxtacellularly labeling single HD cells in subregions of the ADn in awake mice during passive rotation. We identified HD cells that exhibited different response profiles to light, sound, and movement. We also identified a mediolateral gradient of calretinin-expressing (CR+) ADn cells, with CR+ HD cells having narrower tuning widths, lower maximal firing rates, and different intrinsic properties compared to CR-cells. Axons of labeled HD cells could be followed to the retrosplenial cortex, with collaterals innervating the thalamic reticular nucleus (type I cells); others additionally innervated the dorsomedial striatum (type II cells). Most medial CR+ cells preferentially projected to ventral retrohippocampal regions. Surprisingly, we also identified a subpopulation of medial CR+ cells with twisted dendrites and descending axons that avoided the thalamic reticular nucleus, termed tortuosa HD cells (type III cells). We conclude that HD cells of the mouse ADn comprise distinct cell types, providing parallel head-direction-modulated sensorimotor messages to synaptic target neurons within the head direction network.

## Introduction

Spatial navigation is fundamental for survival. Head direction (HD) cells in the mammalian brain are required for spatial orientation, providing a dynamic representation of the position of the head with respect to environmental (allocentric) and body-based (egocentric) cues (Taube, 2007; Alexander *et al*., 2023; Clark *et al*., 2024). In hippocampal and parahippocampal regions of the cerebral cortex, coordinated firing of HD cells with other spatially-modulated neurons, including grid cells, border cells, and place cells, generates a so-called cognitive map of space, which enables spatial navigation based on past experience stored in memory (O’Keefe & Nadel, 1978; Buzsaki & Moser, 2013; Gibson *et al*., 2013; Clark & Harvey, 2016). How is this cognitive map updated? The anterodorsal nucleus of the thalamus (ADn) contains a high density of HD cells, which receive sensorimotor messages primarily via the lateral mammillary nucleus (Taube, 1995; Stackman & Taube, 1998; Taube, 2007). The ADn mainly projects to the granular retrosplenial cortex (RSg) and dorsal presubiculum (PrSd, or postsubiculum), which also contain HD cells (Taube *et al*., 1990; Shibata, 1993a; b; Tukker *et al*., 2015; Hintiryan *et al*., 2025). These signals are then integrated with other spatial signals to update changes in spatial orientation.

The ADn is part of the anterior thalamic nuclear group (ATN), which includes the anterodorsal, anteroventral (AV) and anteromedial (AM) thalamic nuclei. Although these nuclei share some cortical targets (Sripanidkulchai & Wyss, 1986; Shibata, 1993b; a), they are not directly inter-connected, and the AV and AM receive ascending input from the medial rather than the lateral mammillary nucleus (Guillery, 1956; Hayakawa & Zyo, 1989; Vann *et al*., 2007). The AV and AM also contain HD cells, but at a lower density and typically with different temporal patterns compared to the ADn (Tsanov *et al*., 2011; Jankowski *et al*., 2015; Lomi *et al*., 2023; Ji *et al*., 2025). Interestingly, when compared to non-HD cells, ATN HD cells in the mouse increase their firing rate in response to sound stimuli, whisker stimulation, and social touch (Blanco-Hernandez *et al*., 2024). In contrast, the firing of some but not all rat ATN HD cells have also been shown to be strongly suppressed by the experimenter firmly holding and passively rotating the rat (Knierim *et al*., 1995; Taube, 1995). It is not clear whether all ADn HD cells transiently respond to sensorimotor stimuli. Furthermore, based on connectivity, distinct medial and lateral subpopulations of mouse ADn cells have been recently described (Hintiryan *et al*., 2025). This suggests there may be different kinds of HD cells within the ADn providing parallel ‘HD channels’ to the cortical mnemonic system, which would promote more refined models of spatial navigation and memory.

Here we use glass electrode extracellular recordings followed by juxtacellular labeling to define the firing patterns and connectivity of single HD cells in the mouse ADn. We identified three types of projection patterns, a gradient of calretinin (CR) expressing HD cells, and various responses to sensoriomotor stimuli, suggesting the ADn contains ‘parallel channels’ conveying HD-modulated signals to the rest of the HD network.

## Results

### Identification of HD cells in the mouse ADn

We implanted mice with head-plates and lowered a glass electrode into the anterior thalamus during head restraint (Fig. 1A). Once we reached the target depth, we rotated the setup to detect HD cells, which fired strongly in a specific direction, termed the preferred firing direction, or PFD (Fig. 1A, B). After performing extracellular recordings of individual HD cells at different locations bilaterally, we juxtacellularly labeled one HD cell per hemisphere for *post-hoc* recovery in brain sections (Fig. 1C, D, S1, S2). Here we report 94 HD cells (defined by a mean vector length, *r*, of at least 0.3) localized to the ADn (Fig. 1D, E, S1, S3A, Table S1). We recorded HD cells covering the full directional range of the setup (Fig. S3B) and a variety of tuning widths (Fig. S3C). For unidirectional HD cells (n=92/94 cells), the peak firing rate was 23.7 [11.8-42.2] (median [IQR]) Hz (Fig. 1E, S3D), and the background firing rate (i.e. outside the PFD), was 0.78 [0.3-1.7] Hz (Fig. S3E). The directional tuning width was 103.1 [48.1-180]°, with a directional information content (representing how much HD information is conveyed by each spike) of 0.78 [0.53-1.13] bits (Fig. S3C, F). The sparsity (the proportion of the tuning curve that the cell is responsive to) was 0.47 [0.34-0.6], and directional coherence (a measure of smoothness of the HD tuning curve) was 0.75 [0.54-0.87] (Fig. S3G, H). These values are broadly similar to HD cells previously recorded in both head-restrained (Blanco-Hernandez *et al*., 2024) and freely-moving rodents (Taube, 1995; Yoder & Taube, 2009; Clark *et al*., 2024). In addition, we observed burst firing of HD cells in their PFDs (Fig. S3I-N) (Grieves *et al*., 2022; Jiang *et al*., 2024).

**Figure 1.**
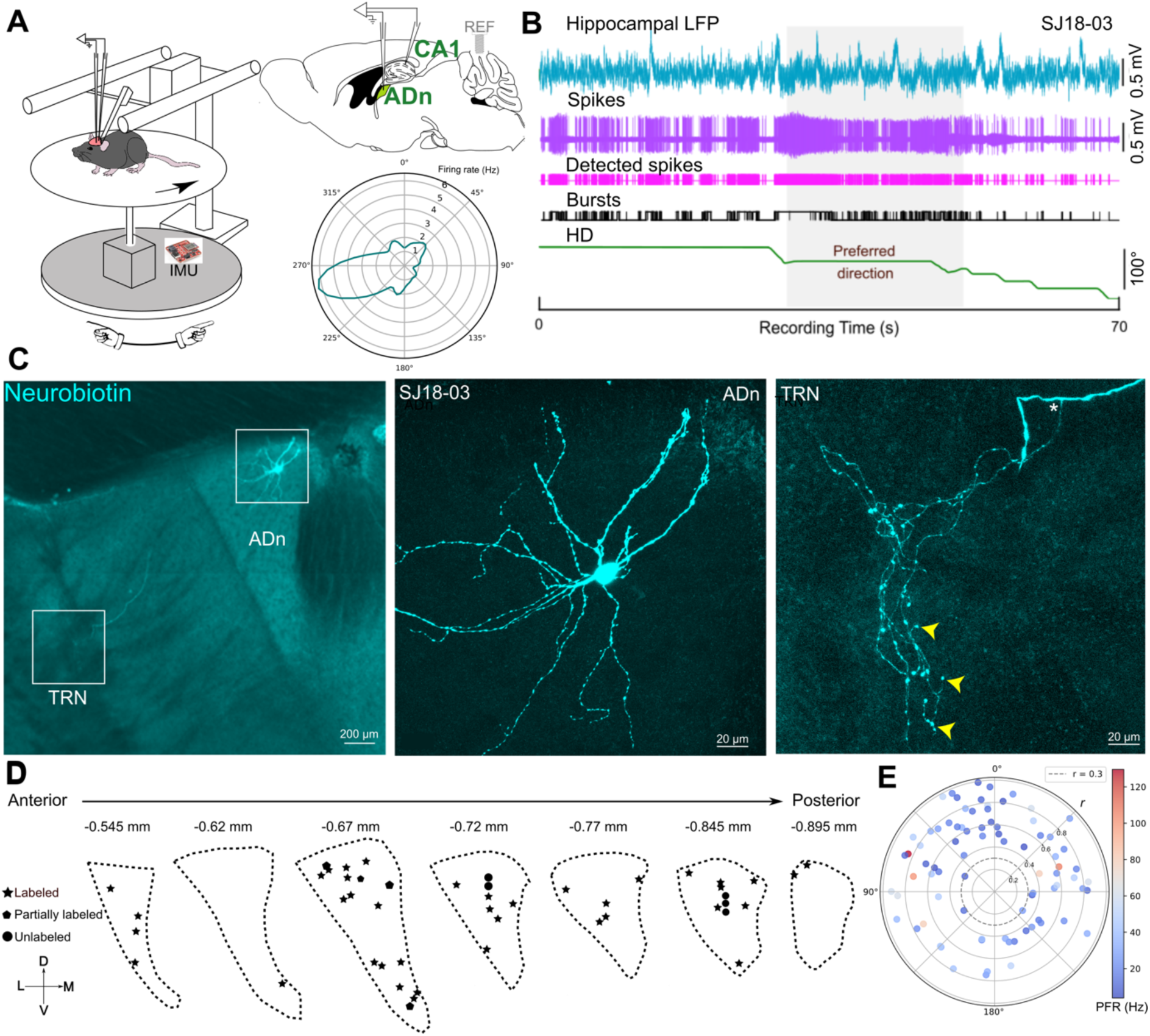
Distribution of single extracellularly recorded ADn HD cells. (**A**) Left, schematic of *in vivo* extracellular recordings in head-restrained mice. The turntable (gray) is passively rotated. An inertial measurement unit (IMU) is used to compute the turning angle. Top right, schematic sagittal brain section indicating target locations of glass electrodes. REF, electrical reference screw above the cerebellum. Bottom right, polar plot of the tuning curve from a recorded ADn HD cell (cell SJ18-03) with its peak firing rate. (**B**) Example simultaneous recordings of the CA1 LFP (cyan), action potentials of the HD cell (SJ18-03, magenta and pink), spike bursts (black), and head direction (HD, green). The shaded area marks the preferred firing direction. (**C**) Left, epifluorescence micrograph of a coronal brain section showing the juxtacellularly neurobiotin-labeled HD cell (cyan, SJ18-03). Middle and right, confocal maximum intensity z-projections (78.3 µm and 74.7 µm thick), showing the soma, partial dendrites, and axon terminals (e.g. arrowheads). Asterisk, main projection axon in the TRN and the origin of the collateral. (**D**) Schematic of different antero-posterior levels of the ADn showing locations of recorded cells (stars, 41 labeled cells; circles, 6 unlabeled cells). See also Extended Data 1 for cell names and Extended Data 2 for representative images. (**E**) Polar plot showing the distribution of the recorded cells. Radial axis, mean vector length (*r*); preferred firing direction, angle; peak firing rate (PFR, color bar, Hz).

**Figure 2.**
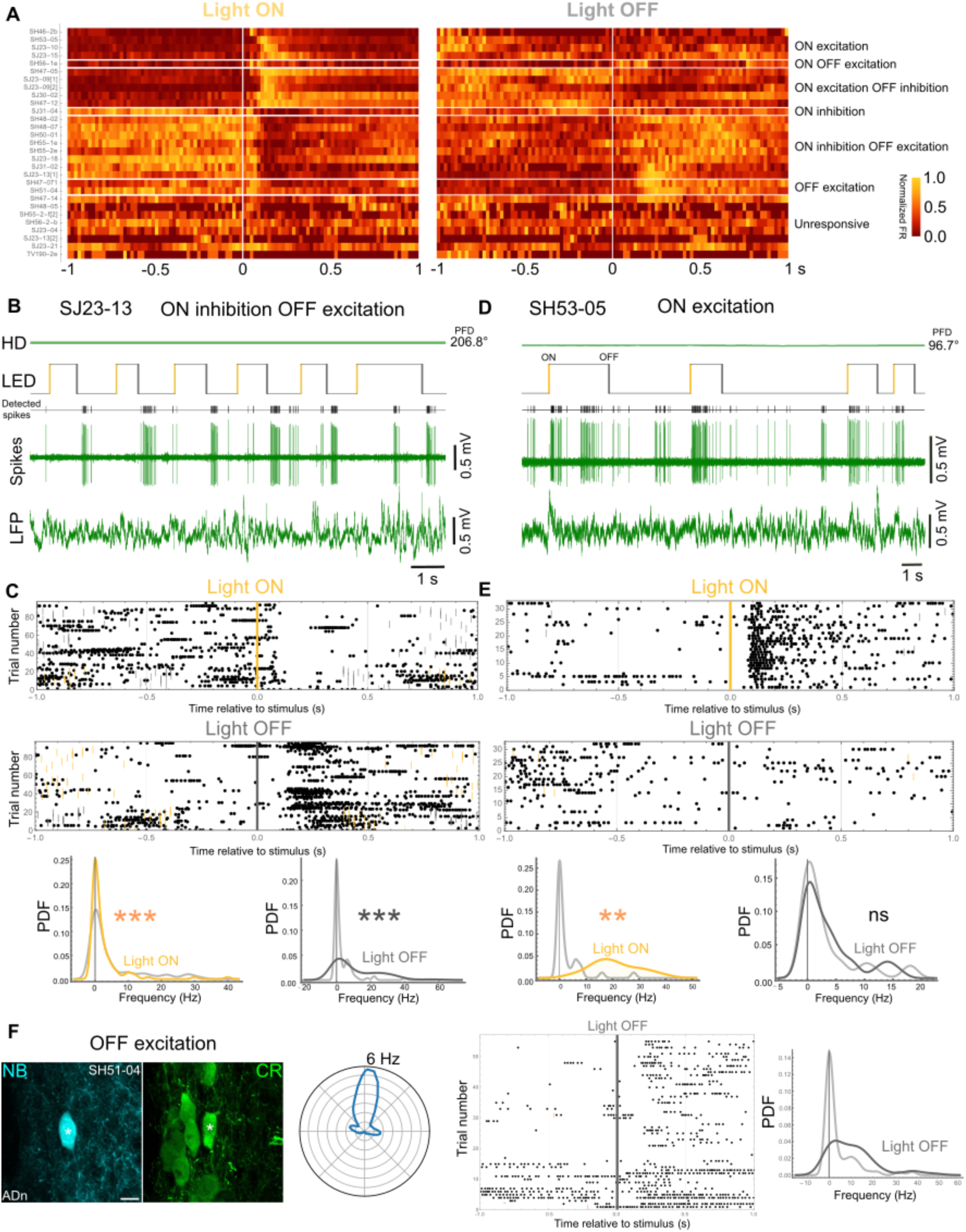
Differential responses of ADn HD cells to light stimuli. (**A**) Heat map of responses of all tested HD cells to light. Bin size, 20 ms. Firing rate (FR) is normalized per row (color bar). Categories (separated by horizontal lines): ON-excitation (n=4), ON-excitation and OFF-excitation (n=1), ON-excitation OFF-inhibition (n=4), ON-inhibition (n=1), ON-inhibition OFF-excitation (n=9), OFF-excitation (n=2), unresponsive (n=8). [1], recorded only in the PFD, or [2], only in the UPFD. (**B**) An HD cell that reduced its firing rate at light ON and increased firing at light OFF (ON inhibition OFF excitation, cell SJ23-13). Top to bottom: head direction (HD, maintained in its PFD), LED stimulation (orange, light ON; dark gray, light OFF), detected spikes, spikes, CA1 LFP. (**C**) Top, raster plots for cell in B. Each point is a spike. Orange ticks, other light ON events; dark gray ticks, other light OFF events. Bottom, probability density functions (PDFs) of firing rates 0.5 s before versus 0.5 s after light ON or OFF. (**D**) An HD cell showing an abrupt and transient increase in firing at light ON but not at light OFF (ON excitation, cell SH53-05). (**E**) Top, raster plots of responses of cell in D. Note consistent latencies for light ON trials. Bottom, PDFs. (**F**) Left, confocal maximum intensity z-projection (10.3 µm thick) of a labeled ADn HD cell (SH51-04, asterisk, neurobiotin, NB, cyan) that was immunopositive for CR (green). Scale bar, 10 µm. Middle, polar plot of HD tuning. Peak firing rate of 6 Hz (radial axis). Right, raster plot and PDF showing the increase in firing after light-OFF (OFF excitation). Not significant, ns.

### ADn HD cells differentially respond to flashes of light

Head direction tuning is dependent on the integration of a variety of sensorimotor-related messages, including visual signals (Zugaro *et al*., 2003; Peyrache *et al*., 2019). Interestingly, the ADn also receives direct input from the retina (Conrad & Stumpf, 1975; Morin & Studholme, 2014). We tested whether ADn HD cells exhibited ‘short-latency’ responses to flashes of bright light (n=27 cells from 12 mice), and observed seven types of responses based on the comparison of average firing rates 500 ms before and after light-ON or light-OFF events (Fig. 2A, Table S2).

The most common response we observed was a transient decrease in firing after light-ON (‘ON inhibition’) followed by an increase in firing after light-OFF (‘OFF excitation’) (n=8/27 cells; Fig. 2A-C). The second largest group increased its firing after light-ON followed by a decrease in firing after light-OFF (‘ON excitation OFF inhibition’; n=4/27 cells; Fig. 2A). We also observed cells that only responded after light-ON (‘ON excitation’; n=4/27 cells; Fig. 2A,D,E), cells that specifically increased their firing rate after light-OFF (‘OFF excitation’; n=2; Fig. 2A,F), decreased their firing rate after light-ON (‘ON inhibition’; n=1; Fig. 2A), and a cell that responded after both light-ON and light-OFF (‘ON OFF excitation’; n=1; Fig. 2A). Responses were specific, as we also found cells that were not significantly modulated by light (i.e. unresponsive to the light stimuli; n=7/27 cells; Fig. 2A, Table S2).

Next we analyzed the response latency and response magnitude (see definition in Methods) for excitation and inhibition. When we compared cells that increased firing after light-ON (ON-excitation, n=10) to cells that increased firing after light-OFF (OFF-excitation, n=11), ON-excitation cells had significantly shorter latencies (90.7 ± 20.2 ms vs 201.2 ± 34.4 ms, \**p* = 0.02, unpaired t-test) and a larger response magnitude (18.5 ± 4.3 Hz vs 5.6 ± 1.1 Hz, \*\**p* = 0.007, unpaired t-test; Fig. S4A). In contrast, for cells that were inhibited after light-ON (ON-inhibition, n=9) versus light-OFF (OFF-excitation, n=5), there were no significant differences for the time to inhibition (95.6 ± 8.1 ms for ON-inhibition vs 76.0 ± 31.9 ms for OFF-inhibition, *p* = 0.687, unpaired t-test; Fig. S4B) or the response magnitude (−11.1 ± 3.5 Hz for ON-inhibition vs −10.2 ± 4.5 Hz for OFF-inhibition, *p* = 0.87, unpaired t-test; Fig. S4B). These data demonstrate the existence of subpopulations of HD cells with different responses to flashes of light.

### Sound and body-movement sensitivity of ADn HD cells

Some HD cells increase their firing rate in response to ‘clicks’ (Blanco-Hernandez *et al*., 2024). We confirmed this by presenting ‘click’ auditory stimuli to 21 ADn HD cells from 10 mice (Fig. 3; Table S3). When the mouse faced the cell’s PFD, 9/17 cells increased their firing rate after the stimulus (‘sound activated’, 15 [10.5-29] ms latency, 23.6 ± 4.2 Hz response magnitude), with 5/9 cells exhibiting a sustained response (maintaining a high firing rate for at least 1 s; Fig. 3A-C). We also observed 1 cell that decreased its firing (Fig. 3D, E); the remaining 7 cells had no significant change in rate. We also observed one HD cell that significantly decreased its firing rate after the stimulus outside the PFD (in the ‘unpreferred’ firing direction, UPFD). Some cells were multimodal, responding to both light and sound stimuli (Table S2 and S3).

**Figure 3.**
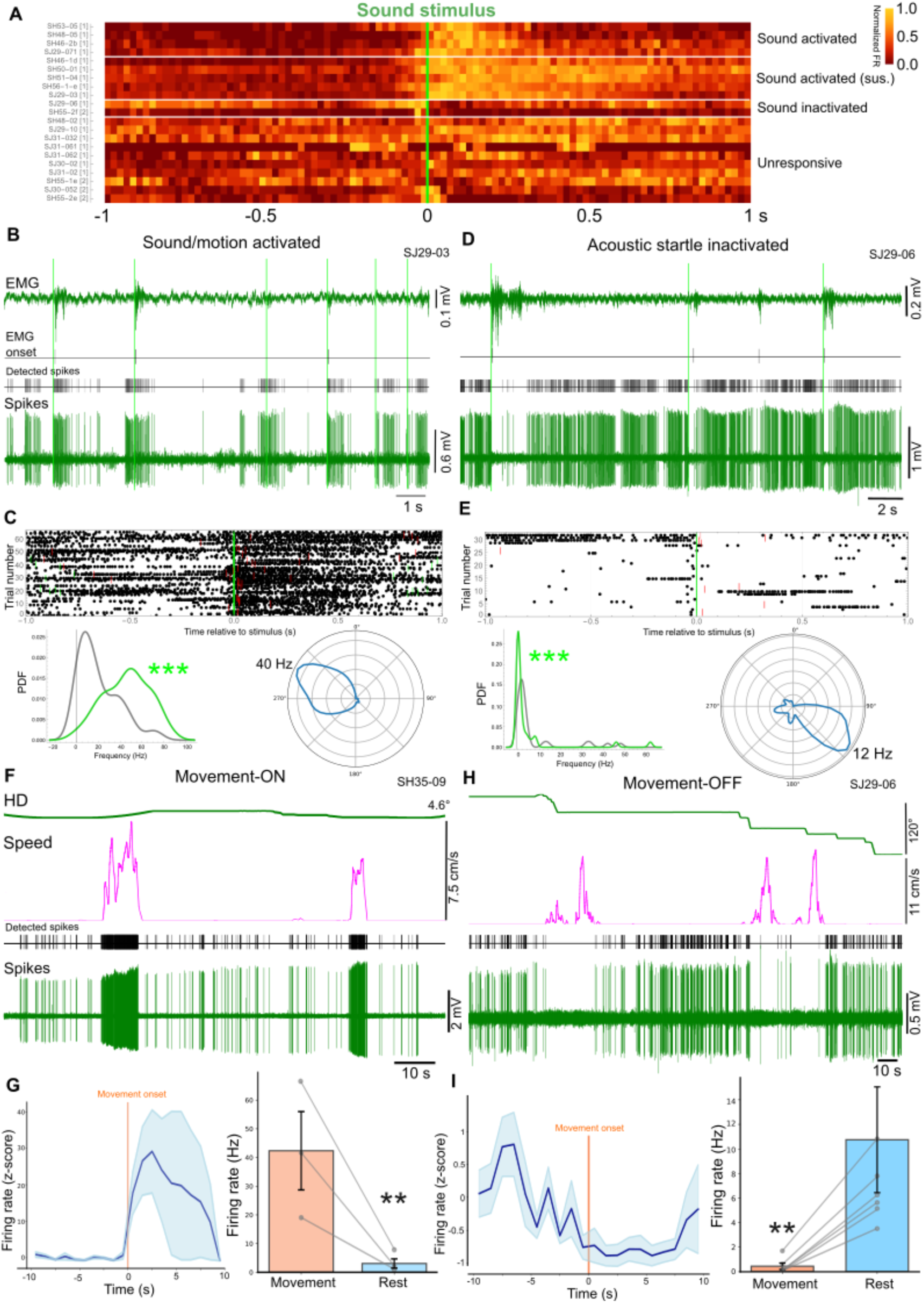
Responses of ADn HD cells to sound and body movement. (**A**) Heat map of responses of all tested HD cells to sound. Bin size, 20 ms. Firing rates are normalized per row (color bar). Categories: sound activated (n=4), sound activated sustained (n=6), sound inactivated (n=2), unresponsive (n=10). [1], recorded only in the PFD, or [2], only in the UPFD. (**B**) An HD cell showing an abrupt and transient increase in firing at sound onset (sound activated, cell SJ29-03). These trials occurred while the animal was resting (running speed = 0), and the cell was in its PFD. ‘Click’ sound stimuli are indicated with green bars. Top to bottom: EMG trace; detected evoked EMG responses; detected spikes; spike trace. (**C**) Top, raster plot of responses of cell in B. Each point is a spike. Green ticks, other sound onset events; red ticks, evoked EMG onset times. Bottom left, PDFs of firing rates 0.5 s before (gray) and 0.5 s after (green) sound onset. Bottom right, polar plot of HD tuning curve with its peak firing rate. (**D**) An HD cell that reduced its firing rate following the sound stimulus coincident with evoked EMG activity (acoustic startle inactivated, cell SJ29-06). (**E**) Raster plot, PDFs, and polar plot with its peak firing rate for cell in D. (**F**) A movement-ON HD cell (SH35-09), showing a large increase in firing specifically during running. From top to bottom: mouse head direction, mouse running speed, detected spikes, spike trace. (**G**) Left, z-scored firing rate of the HD cell in F (normalized based on average firing rate 10 s before movement onset), aligned to the onset of movement (10 s before and after; 1 s time bins). Shaded area indicates ±SEM. Right, bar chart of firing rate during movement and rest (paired data). (**H**) A movement-OFF HD cell (SJ29-06). Note suppression of firing during running. (**I**) Z-scored firing rate and bar chart for cell in H.

We noticed that mice twitched after many of the click stimuli, which we termed the acoustic startle response. To test whether ADn HD cells responded to the sound itself, were evoked by the twitches, or responded to both, we acquired EMG signals simultaneously with single-neuron extracellular recordings (Fig. 3B-E, S5). We split trials into those with EMG responses (i.e. twitches) and those without, and found that HD cell click stimulus responses could be ‘boosted’ by twitches, yet firing rates still significantly increased even on trials without EMG responses (Fig. 3B, C, S5A). We also observed HD cells that only responded when click stimuli were followed by EMG responses, which could either lead to a decrease in firing (acoustic startle-inactivated; Fig. 3D, E, S5B) or an increase in firing (acoustic startle-activated; Fig. S5C).

Next we asked whether more general body movements modulated the firing of HD cells. Indeed, we observed cells that consistently fired at high rates when the mouse spontaneously ran on the running disc (n=11/18, movement-ON HD cells; Fig. 3F, G), and those that abruptly stopped firing during running periods (n=5/18, movement-OFF HD cells; Fig. 3H, I), and unaffected ones (n=2/18). Overall, these data show that ADn HD cells show differential, multimodal responses to sound and body movement.

### Calretinin-expressing HD cells form a medio-lateral gradient in the ADn

We tested juxtacellularly-labeled HD cells for molecular markers that might define HD cell subpopulations, and found that around 30% (n=8/25) of tested HD cells were immunopositive for CR (Table S1, S4, S5). Calb2-expressing cells (for CR) have previously been excluded from molecular classifications of the ADn due to the high Calb2 expression levels in adjacent thalamic nuclei (Kapustina *et al*., 2024). However, we detected CR+ neurons along the medial aspect of the ADn, adjacent to the stria medullaris and clearly within the boundaries of the ADn (defined by the high biotin levels), forming a mediolateral gradient (Fig. 4A-C, S6A, B, S7-S9). Weakly CR+ neurons were intermingled with those lacking detectable immunoreactivity (CR-), with the highest density of CR-neurons at the lateral aspect, bordering the AV and laterodorsal nuclei (LD) (Fig. 4A-D, S6A, S7-S9). The medial band of ADn CR+ cells extended ventrally into the interanterodorsal thalamus (IAD; Fig. S7). We quantified the proportion of CR+ cells within the ADn and found that 27.82% of the total population were CR+ (122 ± 23 CR+ cells/ 472 ± 103 DAPI nuclei from n=3 mice). The mediodorsal (MD) and paraventricular (PVT) nuclei also contained an abundance of CR+ cells (Fig. 4A-D, S7-S9) (Matyas *et al*., 2018). Lateral to the ADn, the LD had dense CR immunoreactivity, whereas the AV lacked CR. This pattern was in stark contrast to PCP4, which marked the AV but not ADn (Fig. S6C).

**Figure 4.**
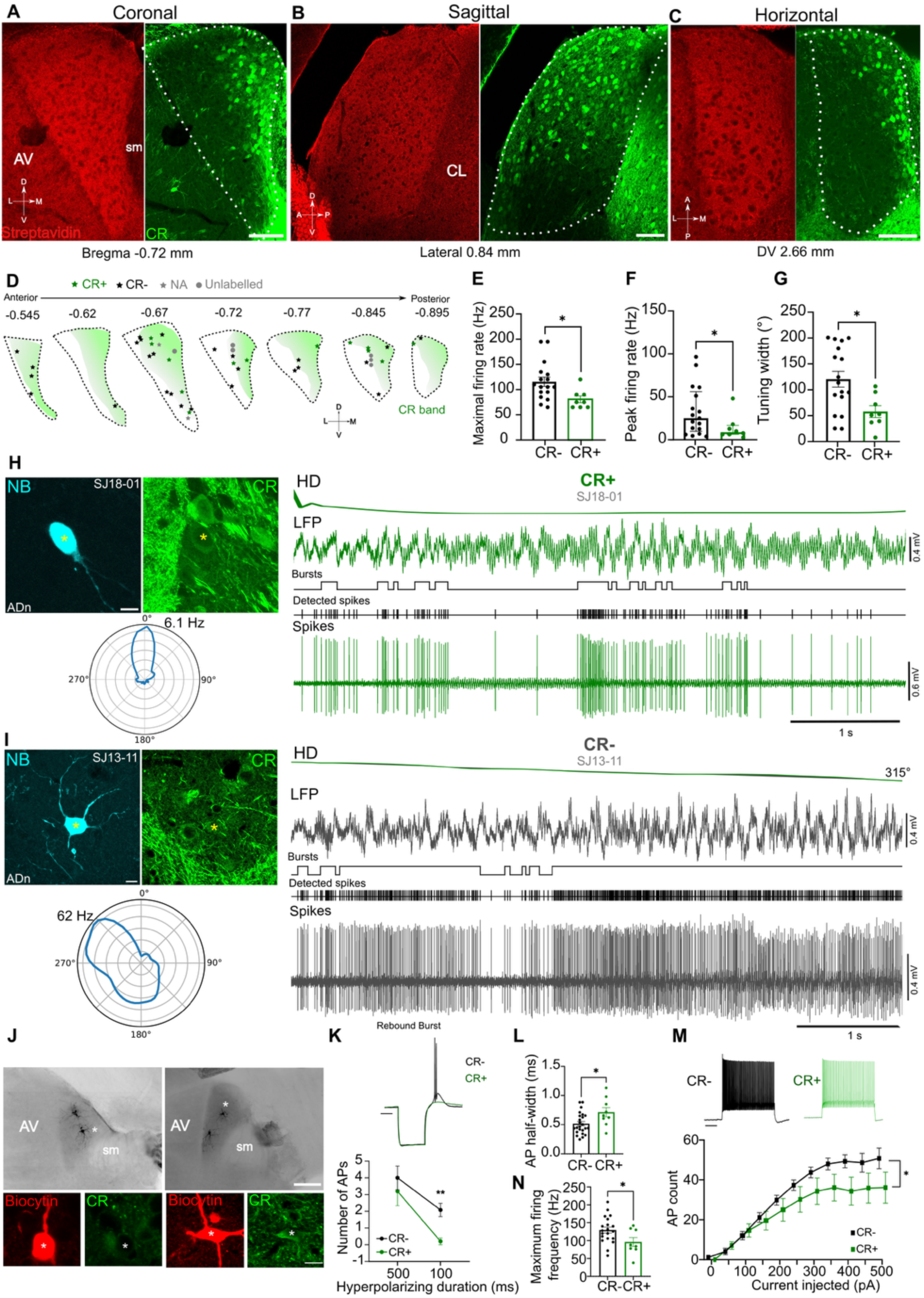
Comparisons of CR+ and CR-ADn cells. (**A**-**C**) The mouse ADn (dashed area) visualized by streptavidin (for biotin, red) at three different orientations. Note the medio-lateral gradient for CR. Confocal single optical sections. Scale bars, 100 μm. DV, dorso-ventral position from brain surface. (**D**) Schematic showing locations of identified ADn cells along with their CR immunoreactivity. Shaded areas indicate the CR zone. NA, CR immunoreactivity not available. (**E-G**) CR+ cells showed lower maximal and peak firing rates and narrower tuning widths compared to CR-cells. (**H**-**I**) Left, a labeled CR+ HD cell (SJ18-01) and a CR-HD cell (SJ13-11) in the ADn (asterisks). Confocal maximum intensity z-projections (3.5 µm and 12.9 µm thick); neurobiotin (NB, cyan); CR (green). Scale bars, 10 µm. Below, polar plots of HD tuning curves with peak firing rates indicated. Right, firing patterns in the PFD. Top to bottom: HD angle, CA1 LFP showing transient theta oscillations, detected bursts, detected spikes, spike traces. Note 50 Hz electrical noise is present in the LFP. (**J**) Examples of biocytin-labeled ADn cells recorded *ex vivo*. Left, the cell denoted by the asterisk was tested for CR (green, inset) and was immunonegative (CR-). Right, a CR+ cell. Biocytin, black (main images), red (insets). Scale bar: 100 μm, inset 5 μm. (**K**) Representative traces of voltage responses following a hyperpolarizing step (−250 pA) from a CR-(black) and a CR+ (green) cell illustrating the lack of rebound burst in CR+ cells. Scale bar: 50 ms. (**L**) Representative traces of voltage responses following a depolarizing step (+500 pA) from CR-(black) and CR+ (green) cells illustrating the decreased firing frequency in CR+ cells. Scale bar: 100 ms. (**M**) Paired-comparison of mean (± SEM) number of APs within a burst at two different durations of current injections. (**N**) Number of action potential (AP) within the rebound burst (n=13 CR- and n=7 CR+ cells from 8 mice). (**O**) AP half-width in ms (n=22 CR- and n=8 CR+ cells from 8 mice). (**P**) The average maximum firing frequency of CR- and CR+ cells. (**Q**) Average AP frequency in response to 0-500 pA current steps illustrating a significant decrease in firing frequency in CR+ cells at high-current injections.

### Calretinin marks two subpopulations of ADn cells

We next mapped the distribution of both labeled (n=40) and unlabeled extracellularly recorded cells (n=8) (Fig. S1; Methods). Based on their distributions in the ADn, we divided them into medial (n=10), middle (n=12) and lateral (n=13) groups (Fig. S6D), as medio-lateral position is known to relate to their preferential target regions (Hintiryan et al 2025). Medial ADn cells had lower maximal firing rates compared to both middle and lateral ADn cells (medial: 67.5 [65-95] Hz, middle: 120 [91.25-128.8] Hz, lateral: 112.5 [88.75-151.3] Hz; Kruskal-Wallis test, \**p* = 0.015; Dunn’s test, medial vs middle: \**p* = 0.03, medial vs lateral: \**p* = 0.02; Fig. S6E), and a lower burst index than middle ADn cells (medial: 0.68 ± 0.05, middle: 0.82 ± 0.03, lateral: 0.7 ± 0.04; one-way ANOVA, *F*_2,32_ = 4.12; Tukey’s test, medial vs middle: \**p* = 0.038; Fig. S6F).

Next, we tested whether CR immunoreactivity was associated with any differences in the firing of ADn HD cells. We found that CR+ cells had significantly lower maximal and peak firing rates (maximal firing rate: 80.6 ± 6.7 Hz [n=8 CR+ HD cells] vs 115.6 ± 9.2 Hz [n=11 CR-HD cells], unpaired t-test, \**p* = 0.022; peak firing rate: 8.8 [6.3-16.7] Hz [CR+] vs 25.1 [9.7-56.1] Hz [CR-], Mann-Whitney test, \**p* = 0.036; Fig. 4D-F; Table S4), consistent with the medio-lateral differences. CR+ cells also had narrower tuning widths compared to CR-cells (57.9 ± 11.5° [CR+] vs 120.6 ± 15.3° [CR-], unpaired t-test, \**p* = 0.015; Fig. 4G; Table S4). Two representative cells are shown in Fig. 4H and I.

To gain insight into why CR+ HD cells had narrower tuning widths and lower firing rates *in vivo*, we examined the intrinsic properties of CR+ and CR-cells in *ex vivo* acute slices using whole-cell patch clamp recordings. We sampled cells in different locations of the ADn to match the *in vivo* locations, and recorded 34 cells of which 8 were CR+ (23.5%) and 22 were CR-(64.7%; the remaining 7 ADn cells were not recovered; Fig. 4J, S10, S11). Cells were clamped at their resting membrane potential and increasing currents were injected to quantify both passive and active properties (Fig. 4K-N, S11). ADn cells typically exhibited a rebound burst following a hyperpolarizing current injection of 100 ms duration (19/20 cells; Fig. 4K, S11A). Yet, when the duration of current injection was increased to 500 ms, the number of AP fired during burst was reduced and we observed a significant decrease in the number of APs firing within a burst in CR+ cells compared to CR-cells (0.2 ± 0.2 [CR+] vs 2.0 ± 0.4 [CR-] APs, n=5 vs n=13 cells from 6 mice, Mann-Whitney test, *p* = 0.010; Fig. 4K, S11A). The resting membrane potential, sag potential, and input resistance were similar for CR+ and CR-neurons (n=8 CR+ and n=22 CR-cells from 8 mice; Fig. S11B-D; Table S4)

Next, we examined active properties and found that CR+ neurons had a broader AP width (0.701 ± 0.05 [CR+] vs 0.530 ± 0.04 [CR-], unpaired t-test, *p* = 0.0293; Fig. 4L). The AP threshold, rheobase, AP amplitude and afterhyperpolarization (AHP) potential were similar across groups (Fig. S11E-H). Interestingly, we observed that CR+ neurons had a lower firing rate with increasing current injections (RM ANOVA, (group x current): F(10, 259) = 1.893, \**p* = 0.046; Fig. 4M), and a lower maximum firing frequency (96.3 ± 12.7 [CR+] vs 130.0 ± 8.2 [CR-], unpaired t-test, \**p* = 0.038; Fig. 4N) consistent with the *in vivo* firing pattern differences. These *in vivo* and *ex vivo* data show that CR+ and CR-represent distinct subpopulations of ADn cells.

### Axonal projections and synaptic targets of ADn HD cells

To make predictions about how these differences in firing patterns might influence postsynaptic targets, such as cortical HD cells, we examined the connectivity of mouse CR+ and CR-ADn neurons (Fig. 5-8). We injected ventral retrohippocampal regions (presubiculum and parasubiculum) with a retrograde-spreading AAV (see Methods) and observed labeled cells primarily in the medial ADn (Fig. 5A, B). The distribution overlapped with CR, and indeed the majority of retrogradely-labeled cells were CR+ (74.31%, n=81/109 retrogradely-labeled GFP+ cells within ADn, from n=3 sections of 1 mouse; Fig. 5B-D). Interestingly, 26.14% CR+ cells lacked C1ql2 immunoreactivity (Fig. 5E). Next we injected a Cre-dependent retrograde-spreading AAV in the dorso-caudal part of the retrosplenial cortex and presubiculum of a C1ql2-Cre mouse. This revealed a clear lateral distribution of retrogradely-labeled GFP+ cells in the ADn (Fig. 5F, G). Only 7.5% were immunopositive for CR (from n=2 sections from 2 mice; Fig 5H, I). These data suggest that a subpopulation of medial CR+ ADn cells mainly lack C1ql2 and preferentially project to ventral retrohippocampal areas, whereas a larger subpopulation of lateral mainly CR-ADn cells likely project to more dorso-caudal cortical areas including dorsal retrosplenial cortex, consistent with a recent report (Hintiryan *et al*., 2025).

**Figure 5.**
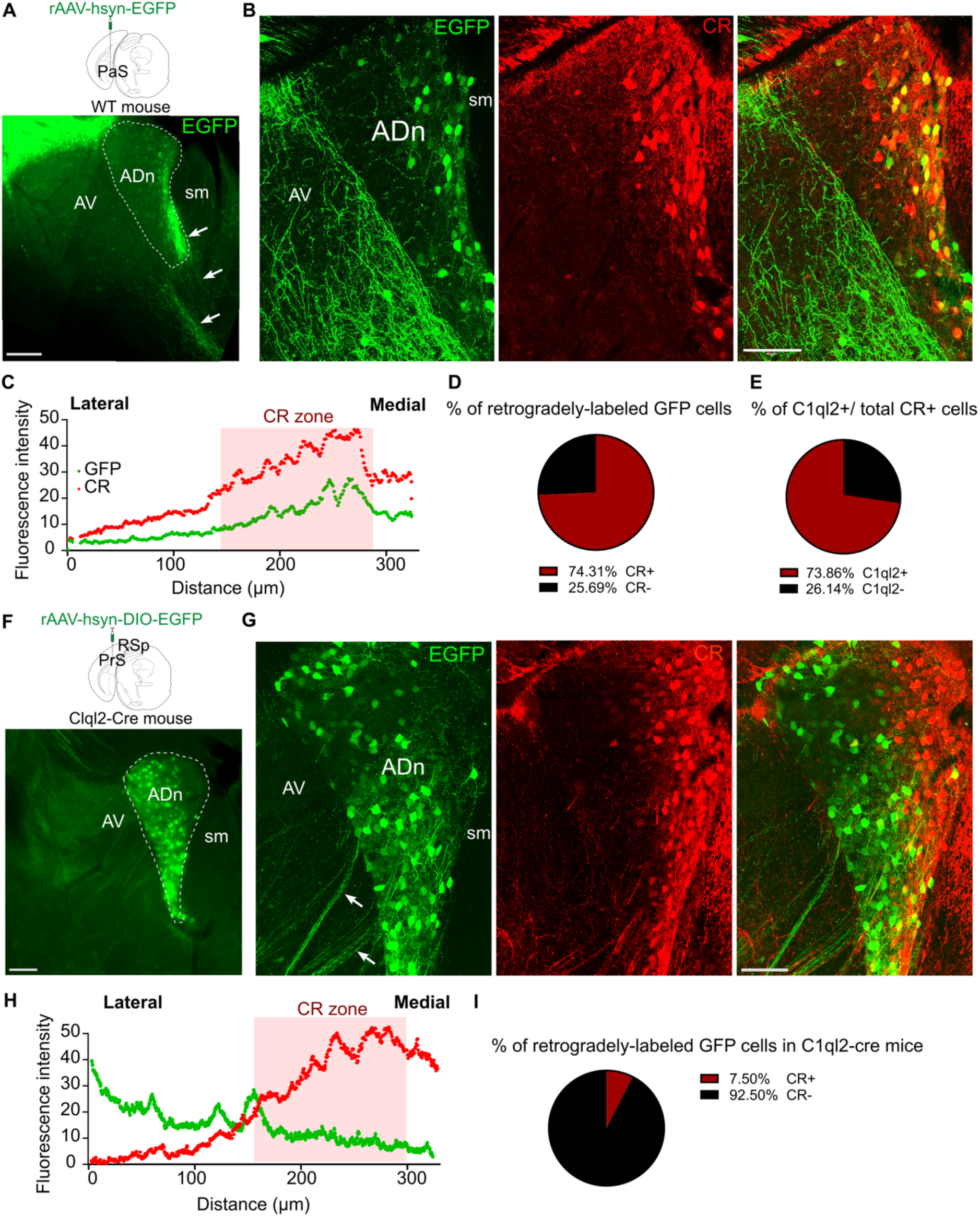
Connectivity of ADn subregions. (**A**) Top, schematic of retrograde AAV (rAAV-hSyn-EGFP) injection into ventral retrohippocampal areas of a wild-type mouse. PaS, parasubiculum. Bottom, widefield epifluorescence of a coronal brain section (case SH63) containing retrogradely labeled cells in the ADn (dashed area) expressing EGFP (green). Note descending axons originating from the medial ADn (e.g. white arrows). Scale bar, 100 µm. (**B**) Retrogradely labeled cells (EGFP+, green) were detected in the ADn and had a similar distribution to CR (red). Case SH63, confocal maximum intensity z-projection (9 µm thick). Scale bar, 100 µm. (**C**) Intensity profiles for GFP and CR. (**D**) Pie chart illustrating the proportion of CR+ cells within the total population of GFP+ cells. (**E**) Pie chart illustrating the proportion of C1ql2+ cells within the total population of CR+ cells. (**F**) Top, schematic of a Cre-dependent retrograde AAV (rAAV-hSyn-DIO-EGFP) injection into the dorso-caudal region of the retrosplenial cortex (RSp) and presubiculum (PrS) of a C1ql2-Cre mouse. Bottom, widefield epifluorescence of a coronal brain section showing selective labeling in the ADn (dashed area), case SH85. Scale bar, 100 µm. (**G**) Retrogradely labeled cells (EGFP+, green) were detected in the ADn and had very little colocalization with CR (red). Note axons originating from the ADn heading laterally towards the TRN (e.g. white arrows). Case SH63, confocal maximum intensity z-projection (9 µm thick). Scale bar, 100 µm. (**H**) Intensity profiles for GFP and CR. (**I**) Pie chart illustrating the proportion of CR+ cells within the total population of GFP+ cells.

Next, we examined labeled ADn cells by following their axons and identifying collaterals (n=13/40 labeled cells; Table S5). Most identified cells were distributed in the middle and lateral regions of the ADn (n=11/13) and followed a specific route from the ADn that we designated as a ‘type I’ projection (n=8/13 cells; Fig. 8H). A representative type I CR-cell (SJ19-02) is shown in Fig. 6. This cell had a higher-than-average maximal firing rate (120 Hz), a high mean vector length (*r* = 0.84; Fig. 6A; Table S1), and was located in the lateral ADn (Fig. 6 B-D). Its dendrites exhibited a relatively complex looping pattern; some distal dendritic tips extended across the border of the ADn into the internal medullary lamina (Fig. 6C). The axon emerged from the medial side of the soma and formed a hairpin turn. It looped back across the soma towards the lateral border, then crossed the AV to reach the TRN where it formed 5 collaterals with terminals (n=109 large terminals observed; Fig. 6C-E). Terminals of SJ19-02, and another type I CR-HD cell SJ18-03, were double immunopositive for vGLUT1 and vGLUT2 (Fig. 1C, 6F, G).

**Figure 6.**
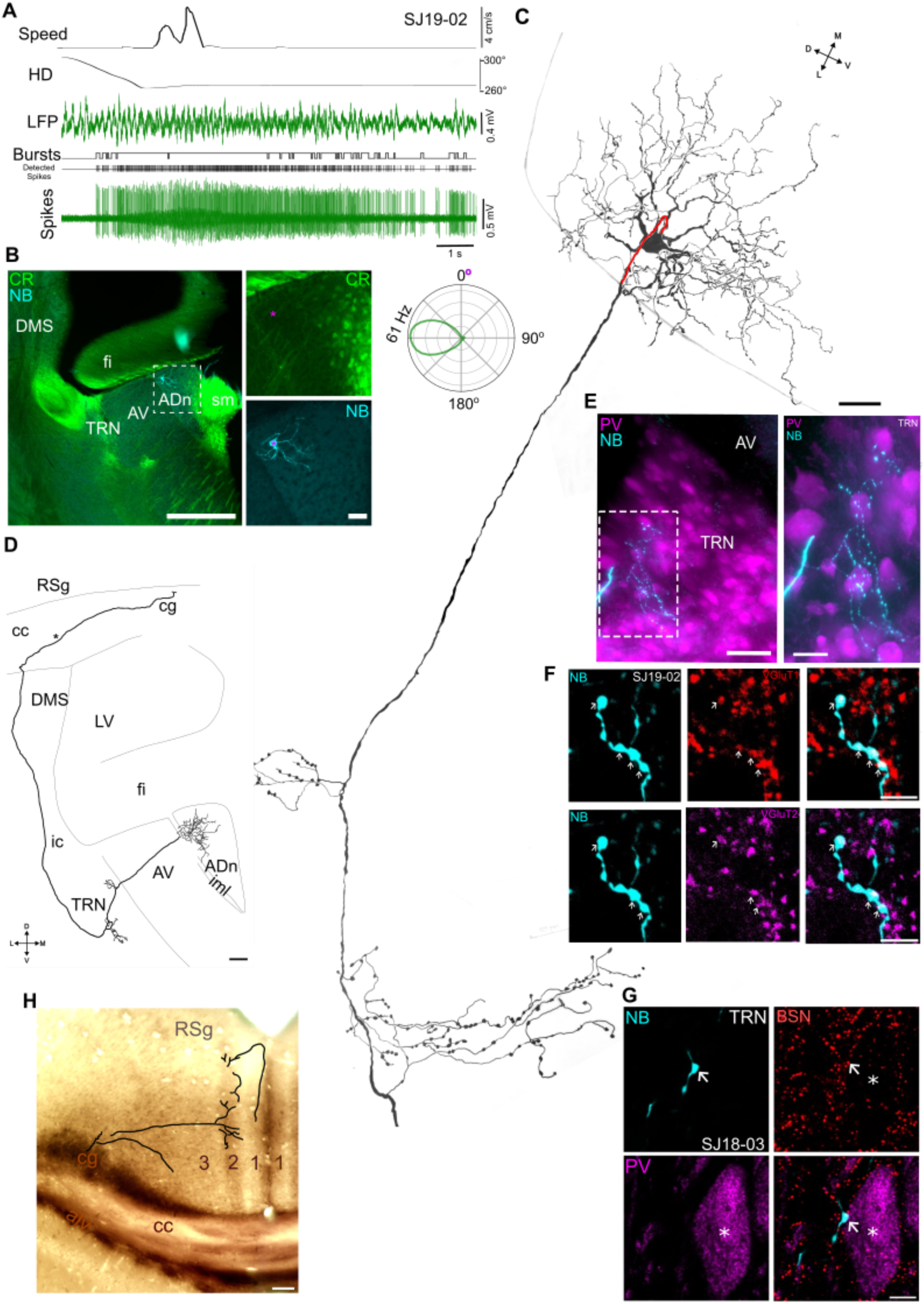
Axon projections and synaptic target regions of ADn HD cells. (**A**) Firing patterns of a labeled HD cell (SJ19-02). Note increase in firing during movement, coincident with CA1 theta oscillations (LFP). Bottom right, polar plot of HD tuning curve with its peak firing rate indicated. (**B**) Left, widefield epifluorescence micrograph of a 70-µm-thick coronal brain section showing part of the soma and dendrites of cell SJ19-02 (asterisk, NB, neurobiotin, cyan) in the lateral ADn. Right, enlarged view of boxed region showing the soma (cyan, asterisk), which lacks detectable CR immunoreactivity (green). (**C**) Full reconstruction of the soma and dendrites of cell SJ19-02 and axon collaterals in the TRN (only large terminals are shown). The axon originates from the medial side of the soma (highlighted red). Reconstructed from 5 70-µm-thick consecutive brain sections. (**D**) A two-dimensional representation of the axon projection of SJ19-02 between the ADn and cingulum bundle (cg) (covering ∼-0.10 to −0.58 mm AP). Asterisk, turning point of the axon. The horizontal bar indicates the end of the reconstructed portion. (**E**) Left, maximum intensity z-projection (37.8 µm thick, widefield epifluorescence) of an axon collateral in the TRN from cell SJ19-02 (neurobiotin, NB, cyan) within the PV-enriched TRN (magenta). Note lack of PV cells in the adjacent AV. Right, enlarged view of box region, maximum intensity z-projection (46.5 µm thick). (**F**) Boutons of CR-HD cell SJ18-03 in the TRN (neurobiotin, NB, cyan, arrows) were immunopositive for both VGLUT1+ (red) and VGLUT2+ (magenta). Confocal maximum intensity z-projection, 1.8 µm thick. (**G**) A PV+ neuron (magenta) in the TRN is innervated by boutons (cyan) of an CR-HD cell (SJ18-03). Bassoon (red) indicates the expected location of synapses, single optical confocal image. (**H**) A two-dimensional representation of the axon collaterals of SJ19-02 in the RSg originating from the main axon in the cg (partial reconstruction from 4 70-µm-thick consecutive brain sections overlaid on one representative section). Note highly branched pattern in layer 3, with axon reaching layer 2 as well as horizontally along layer 1 (axon terminals not shown). Abbreviations: alv., alveus; cc, corpus callosum; LV, lateral ventricle; fi, fimbria; ic, internal capsule; iml, internal medullary lamina. Scale bars (µm): B (left) *500*; D, H *100*; B (right), C, E (left) *50*; E (right), F, G, *5*.

The main axon of cell SJ19-02 continued rostrally via the internal capsule and dorsomedial striatum (DMS); no collaterals were observed in the DMS (Fig. 6D). The axon traversed the corpus callosum and entered the cingulum bundle). At the level of the triangular septal nucleus (∼0.35 mm posterior of Bregma), the axon turned and headed caudally within the medial part of the cingulum. The main axon formed a collateral at the level of the dorsal hippocampus (∼1.35 mm posterior of Bregma) and innervated the RSg. Terminals were observed in layers 1, 2 and 3 (Fig. 6H). The main axon continued caudally in the cingulum and was last observed at the level of the angular bundle (∼ 3.10 mm posterior of Bregma), likely heading to the PrSd. In summary, type I cells projected across the AV and formed collaterals in the TRN, then traveled via the internal capsule and DMS to enter the cingulum bundle before innervating the RSg and other cortical areas (Fig. 8H).

We identified 3 other CR-HD cells that initially followed the same trajectory as the type I cells but formed additional collaterals in the DMS (type II cells; Fig. 8H; Table S5). A representative type II CR-HD cell, SH79-07, was recorded in a similar location to SJ19-02 in the lateral ADn (Fig. 7A-D, S1). A collateral of SH79-07 in the DMS (Fig. 7E) gave rise to terminals that innervated cholinergic neurons (choline acetyltransferase, ChAT+; Fig. 7F). The main axon continued into the cingulum bundle and branched extensively in the RSg (Fig. 7A; collaterals not shown). Therefore, type II cells are distinguished from type I cells by the additional branching in the DMS (Fig. 8H).

**Figure 7.**
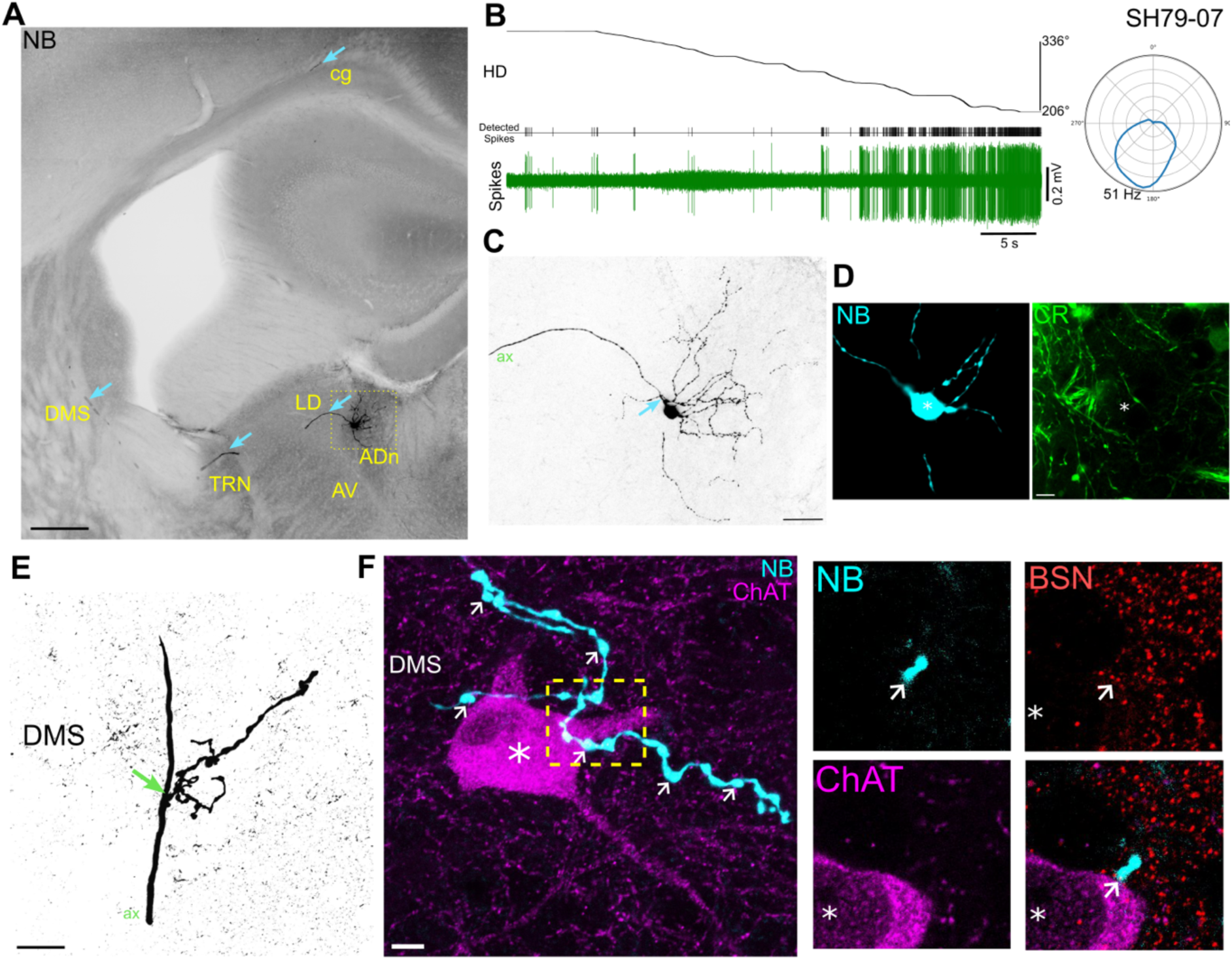
Firing patterns and synaptic targets of a CR-HD cell. (**A**) Widefield epifluorescence micrograph (reverse contrast) of a coronal brain section showing the soma and partial dendritic tree of cell SH79-07 in the lateral ADn, along with the axon (e.g. cyan arrows) projecting to the TRN, DMS, and RSg (neurobiotin, black), forming collaterals in each area (not visible). Scale bar, 300 µm. (**B**) Left, firing patterns of the labeled HD cell (SH79-09) at different angles. Right, polar plot of HD tuning curve with its peak firing rate indicated. (**C**) Enlarged view of soma and dendrites of cell SH79-07 showing the origin of the axon (arrow). Confocal maximum intensity z-projection (51.2 µm thick). Scale bar, 50 µm. (**D**) The soma of cell SH79-07 (cyan, asterisk) lacks CR immunoreactivity (green). Confocal maximum intensity z-projection (6 µm thick). Scale bar, 10 µm. (**E**) Axon collateral (arrow, branch point) arising from the main axon (ax) of cell SH79-07 in the DMS. Confocal maximum intensity z-projection (10 µm thick). Scale bar, 5 µm. (**F**) Left, terminals from a collateral of cell SH79-07 (neurobiotin, NB, cyan, e.g. arrows) near a ChAT+ neuron (asterisk, magenta) in the DMS. Right, single optical section showing innervation of the ChAT neuron by a bouton (NB, arrow) based on bassoon (BSN) puncta (red). Scale bars, 5 µm.

**Figure 8.**
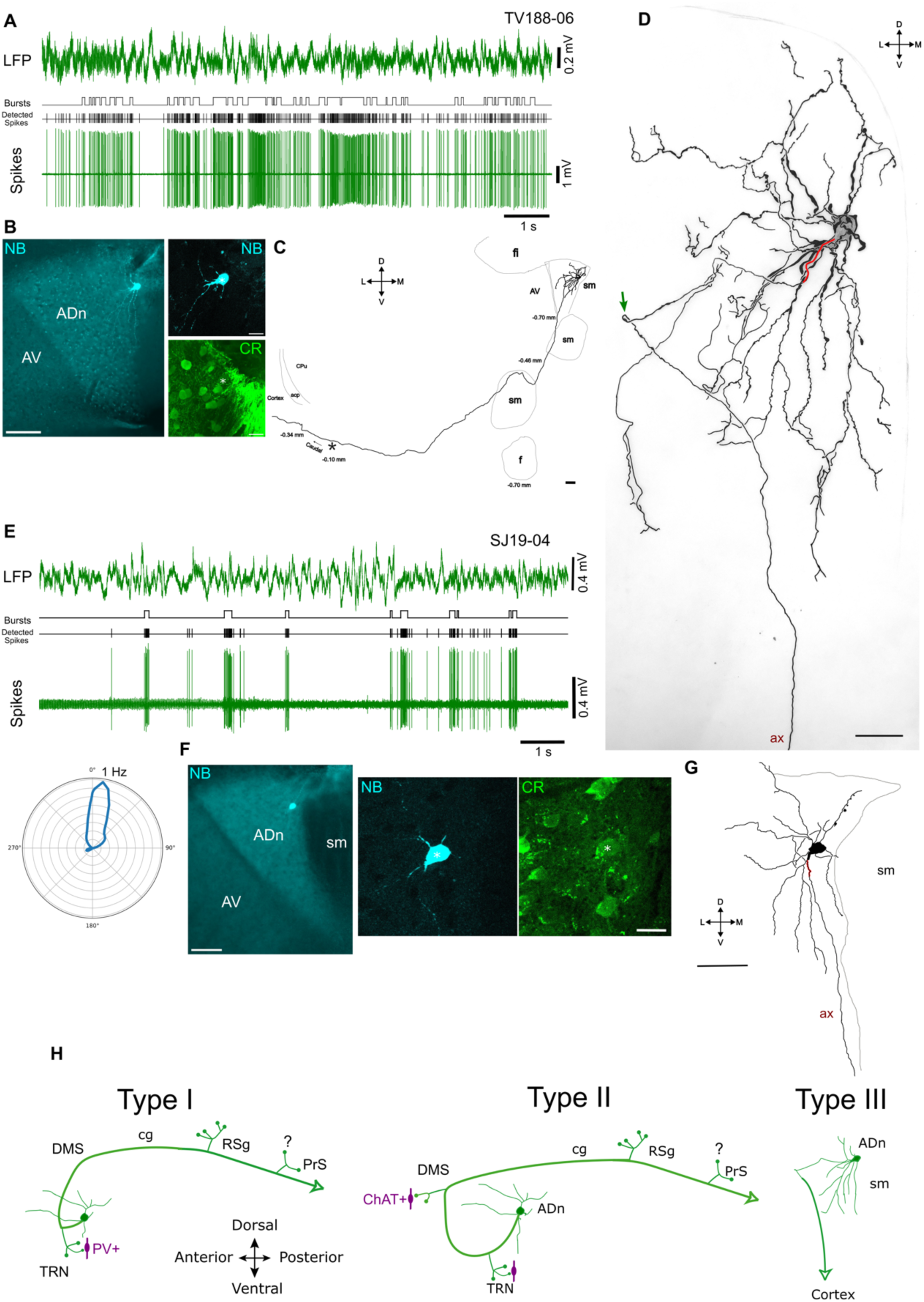
‘Tortuosa’ HD cells are CR+ and have descending axons. (**A**) Firing patterns of a labeled ADn cell (TV188-06) in its PFD. (**B**) Left, widefield epifluorescence micrograph of a coronal brain section showing the soma and partial dendritic tree of cell TV188-06 (asterisk, NB, neurobiotin, cyan) in the medial ADn. Right, the cell was immunopositive for CR (green, asterisk). (**C**) A two-dimensional representation of the axon projection of TV188-06 between the ADn and endopiriform cortex (covering ∼-0.10 to −0.70 mm AP). Asterisk, turning point of the axon. (**D**) 2D reconstruction of the soma and dendrites of TV188-06 (from 4 70-µm-thick consecutive sections). The axon originated from the soma (highlighted red). Arrow, looping point of the axon where it changes direction. The soma is opaque in order to visualize dendrites. Note some dendritic segments were left unconnected due to weak labeling in some sections. (**E**) Firing patterns of a labeled ADn HD cell (SJ19-04) in its PFD. Bottom left, polar plot of HD tuning curve with peak firing rate indicated. (**F**) Left, widefield epifluorescence micrograph of a coronal brain section showing the soma and partial dendritic tree of cell SJ19-04 (asterisk, NB, neurobiotin, cyan) in the medial ADn. Right, the cell was immunopositive for CR (green, asterisk). (**G**) Partial 2D reconstruction of the soma and dendrites from 1 70-µm-thick section. The axon originated from a dendrite (highlighted red) and headed ventrally. (**H**) Schematic summary of three types of ADn cell projections. Green circles represent the axon terminals observed. No terminals of labeled cells were observed in PrS due to limited labeling strength (question mark). Synaptic target neurons (magenta) are shown in apposition to axon terminals in the TRN and DMS. Abbreviations: acp, anterior commissure, posterior; AV, anteroventral thalamic nuclei; ax, axon; cg, cingulum; CPU, caudate putamen (striatum); f, fornix; fi, fimbria; sm, stria medullaris. Scale bars (µm): B (left), C, F (left) 100; B (right), F (right) 20; D, G 50.

We also recorded and labeled two unusual cells in the dorso-medial region of the ADn (TV188-06 and SJ19-04; Fig. 8). Both cells were CR+ (Fig. 8B, F) and fired in bursts within their PFDs (Fig. 8A, E; Table S1). Cell SJ19-04 showed typical CR+ HD tuning with a low peak firing rate and narrow tuning width (Fig. 8E; IMU data for TV188-06 were unavailable). The contorted dendrites resembled the twisted branches of *Salix babylonica var. pekinensis ‘Tortuosa’* (corkscrew willow tree), and some dendrites were found to wrap around other dendrites from the same cell (Fig. 8D). Surprisingly, the axons of both cells headed ventrally from the ADn and did not send collaterals to the TRN (type III cells; Fig. 8C, D, G). The axon of TV188-06 emerged from the ventral part of the soma and headed laterally across the ADn, as if it was heading towards the TRN via AV. However, compared to the majority of HD cells (e.g. Fig. 1C, 6D, E, 7A, C), the axon of TV188-06 sharply looped back over itself and headed ventrally, to leave the ADn close to the stria medullaris (Fig. 8C, D). The axon traveled in the antero-ventral direction along the edge of the stria medullaris. It headed laterally until ∼-0.10 mm posterior of Bregma where it started to head in a latero-posterior direction, traveling ventral of the posterior part of the anterior commissure. It was last observed within the endopiriform cortex (∼0.34 mm posterior of Bregma), still heading in a posterior direction (Fig. 8C). Similar descending axons were also observed from medial (mainly CR+) neurons that were retrogradely labeled from ventral retrohippocampal regions (Fig. 5A-D) but were not observed from the mainly CR-neurons that were retrogradely labeled from the dorso-caudal parahippocampal areas (Fig. 5F-I). Based on the unique descending axons, CR immunopositivity, their location in the dorso-medial ADn, and their distinct twisted dendrites, we name these type III cells as tortuosa HD cells.

## Discussion

Glass electrode extracellular recordings in the ADn of awake head-fixed mice during passive rotation provided us with reliable measures of HD cell activity (e.g. directional tuning, burstiness, sparsity, coherence), which were similar to freely-moving conditions (Yoder & Taube, 2009). Juxtacellular labeling of recorded cells enabled us to define, to our knowledge for the first time, different types of thalamic HD cells in terms of their neuronal activity, neurochemical profile, and axon projections. We observed individual HD cells responding to flashes of light with 6 different patterns, whereas others remained unresponsive. A subset of cells responded to sound stimuli within their receptive fields, and firing could also be modulated by both small and large body movements. Interestingly, some cells were multimodal, responding to both light flashes and sound. We identified both CR+ and CR-cells, which followed a medio-lateral gradient. The CR+ cells generally had lower maximal and peak firing rates and narrower tuning widths compared to CR-cells, which were related to differences in their intrinsic properties. Lastly, we could identify cells with different projections: type I cells followed the ‘typical’ route via the TRN and DMS to enter the cingulum bundle; type II cells formed additional collaterals in the DMS; and type III cells avoided the TRN and had descending axons.

### HD cells as parallel channels

We observed HD cells responding robustly to flashes of light, with a range of ON-OFF response profiles resembling the parallel channels conveyed by ganglion cells of the retina (Roska & Werblin, 2001). For example, ‘ON alpha’ ganglion cells exhibit sustained excitation when a white spot is presented to the retina (i.e. at light ON), ‘OFF alpha’ cells respond transiently when a black spot is presented (i.e. light OFF), and ‘OFF alpha sustained’ cells have sustained responses at light OFF (Farrow *et al*., 2013). The response latencies of ADn HD cells were in the same range of the light responses of cells recorded in the dorsal lateral geniculate nucleus of isoflurane-anesthetized mice (Piscopo *et al*., 2013). Given these similar latencies and that the ADn receives direct input from the retina (Conrad & Stumpf, 1975; Morin & Studholme, 2014), we hypothesize that the ON and OFF excitation responses are directly driven by monosynaptic inputs from different types of ganglion cell. The distribution of these potential inputs remains to be elucidated, but we expect they will map on to specific types of HD cells. The source of the inhibitory responses (i.e. ON or OFF inhibition) is unlikely to be due to a direct inhibitory input from the retina, as most or all ganglion cells are considered glutamatergic (Mimura *et al*., 2002; Gong *et al*., 2006).

Many types of ganglion cells are intrinsically photosensitive due to the expression of opsins such as melanopsin. Opn5-expressing retinal ganglion cells have been shown to project to a specific zone close to the medial habenula (Fernandez *et al*., 2018), which we suggest is the CR-enriched medial ADn. This suggests that in addition to the fast responses to flashes of light, medial HD cells may also be modulated by gradual changes in luminance via intrinsically photosensitive ganglion cells. The rapid responses we recorded suggest HD cells are likely encoding fast global changes to the visual scene, which could help anchor allocentric cues for the HD network.

As well as light responses, many (but not all) HD cells responded to click stimuli, with the majority exhibiting an increase in firing rate. As other HD cells were unresponsive, even in the same animal as responding cells, sound stimuli are likely conveyed via specific cell types with different connectivity to other HD cells. ADn cells have previously been shown to respond to click stimuli even when the mouse is anesthetized (Blanco-Hernandez *et al*., 2024), suggesting that responses can be directly related to sounds. Moreover, we observed that some cells were additionally modulated during the brief muscle twitches caused by the stimulus (cross-modal summation), which is reminiscent of the startle response that activates the vestibular system (Yeomans *et al*., 2002). This suggests that HD cells in the ADn integrate a range of sensorimotor inputs via different pathways (Blanco-Hernandez *et al*., 2024), supporting cue-based realignment when navigating dynamic environments. In contrast, the non-responsive HD cells may contribute to the maintenance of a stable direction signal in the absence of external inputs, possibly relying more heavily on idiothetic information and internal network dynamics.

We also observed that firing rates of different HD cells were modulated by body movement. Taube (1995) reported a weak positive correlation between the rat’s linear speed and ADn HD cell firing rates. We detected HD cells in the mouse that increased, decreased or had no change in their firing rate during locomotion, suggesting that some HD cells may be modulated by running speed. Speed cells have been described in other brain regions, including the medial entorhinal cortex (MEC) (Kropff *et al*., 2015; Hinman *et al*., 2016). In the MEC, positively and negatively speed-modulated subpopulations of HD cells were found (Hinman *et al*., 2016), which may depend on different kinds of speed-modulated HD cells of the ADn; other speed cells fire independently of the HD, suggesting different thalamocortical/corticocortical connectivity (Kropff *et al*., 2015). There are also subpopulations of HD cells in the AV, MEC, and parasubiculum that are modulated by 5-12 Hz theta oscillations, which have been proposed to encode an internal sense of direction (Tsanov *et al*., 2011; Brandon *et al*., 2013; Vollan *et al*., 2025). ADn HD cells are not theta modulated (Taube, 1995) but likely provide strong transient HD-modulated sensorimotor signals to the cortical HD network, which could contribute to the burst-firing of HD-modulated grid cells (Vollan *et al*., 2025). Future studies are needed to elucidate the relative contributions of each ‘HD cell channel’ with respect to the various sensory triggers and how they update the cortical mnemonic system.

### HD cell subpopulations defined by CR

By combining juxtacellular labeling with immunohistochemistry, we found that HD cells in the medial ADn frequently expressed CR, whereas lateral cells were predominantly CR-. The CR+ neurons in the medial ADn are part of a stream of CR+ neurons extending along the midline of the mouse and human thalamus (Matyas *et al*., 2018; Viena *et al*., 2021; Sárkány *et al*., 2024). In general, compared to CR-cells, CR+ HD cells had narrower directional tuning and lower firing rates, and produced fewer spikes per rebound burst. This is in line with other midline thalamic CR+ neurons recorded in anesthetized mice (Lara-Vásquez *et al*., 2016). In the midline paraventricular thalamic nucleus, CR+ neurons respond to transient changes in muscle tone, linked to arousal from sleep (Matyas *et al*., 2018). Although we only labeled one CR+ neuron that was activated by sound, we predict that medial/CR+ ADn cells would be the most sensitive to changes in arousal, such as during behavioral state transitions, responding similarly to what we observed for some cells following muscle twitches induced by the click stimulus. Likewise, other midline CR+ may be sensitive to acoustic startle responses.

In terms of projections, we found that the CR+ subpopulation preferentially targeted ventral retrohippocampal areas (e.g., presubiculum and parasubiculum), whereas CR-cells innervated more dorsal regions including the dorsal part of the retrosplenial cortex. This connectivity pattern fits well with Hintiryan *et al*. (2025), who showed that ADn subdivisions are defined by projection target, supporting the idea of parallel HD streams conveying complementary sensorimotor signals.

To our knowledge, CR is the first marker that has been used to define two subpopulations of HD cells in the thalamus. Kapustina *et al*. (2024) used single-cell RNA sequencing and spatial transcriptomics to define ATN subpopulations but excluded all Calb2-expressing cells from the ADn. Here we unequivocally show that extracellularly recorded and juxtacellularly labeled ADn cells can indeed be CR+, which represented 32% of labeled HD cells.

In the human thalamus, CR+ ADn cells are preferentially vulnerable to tau pathology (Sárkány *et al*., 2024), a hallmark of Alzheimer’s disease and other progressive neurodegenerative diseases. This raises the possibility that the gradual loss of CR+ HD cells during disease progression preferentially affects neural circuits of CR+ cells and their associated functions and firing properties.

### HD cell types and connectivity

Calretinin expression alone is insufficient to define cell types. Axon projections and their targets have proved a reliable means to define cell types in the hippocampus and other cortical areas, especially in combination with firing patterns and neurochemical profiles. We identified 3 different patterns of axonal projections, which we refer to as types I, II, and III. Type I cells were mainly CR- and formed collaterals in the TRN and cortex. Type II cells were CR- and additionally innervated neurons in the DMS. Type III cells were CR+, located in the dorso-medial ADn, and had a completely different projection pattern. We were unable to follow axons to their ultimate target regions due to labeling strength or other technical reasons. Future recordings of single ADn cells in combination with full labeling of the axons will likely reveal additional projection patterns as well as identification of the major postsynaptic target regions.

Two ADn cells have previously been reconstructed (excluding the axon terminals) as part of the MouseLight project (Winnubst *et al*., 2019); one is located in the middle of the dorsal ADn (neuron AA0073) and the other appears to be along the border of the internal medullary lamina in the ventral ADn (neuron AA1445). Both neurons form collaterals in the TRN and project via the DMS. Neither cell forms collaterals in the DMS, resembling our ‘type I’ cells. Neuron AA0073 enters the cingulum bundle and appears to minimally innervate the RSg at 3 or 4 separate antero-posterior levels. The main target is the PrSd but other minor collaterals appear to innervate a small area close to the angular bundle as well as some branching in the parasubiculum. Two branches (without further collaterals) appear to reach the entorhinal cortex, consistent with tracing studies in rat (Shibata, 1993a). The other neuron shows extensive innervation of the entire antero-posterior extent of the RSg, spanning superficial cortical layers, likely resembling the pattern of type I HD cell SJ19-02. The axon of neuron AA1445 does not appear to project further than the RSg. The projection patterns of type I and type II ADn cells we describe, as well as the two from Winnubst *et al*. (2019) share features of the so-called ‘multifocal’ thalamocortical neurons (Clasca, 2023), although it is evident that individual ADn cells can exhibit a range of projection patterns, which reflects their related but distinct target regions (Hintiryan *et al*., 2025).

The type II ADn HD cells, which formed collaterals in the DMS, are likely responsible for the HD signals in the DMS, as lesions of the ATN eliminate DMS HD signals (Mehlman *et al*., 2019b). Tracing studies have also confirmed that a subpopulation of ADn cells innervate the DMS (Mehlman *et al*., 2019a). We detected presynaptic terminals from ADn cells in close apposition to cholinergic neurons in the DMS. Striatal cholinergic interneurons modulate local circuits, suggesting that ADn HD cells may also provide indirect influence on striatal circuits. For example, the type II cells could also contribute to the egocentric boundary responses observed in the DMS (Hinman *et al*., 2019) or could even interact with egocentric boundary responses to help guide movement away from barriers.

The type III cells were unusual in that they avoided the TRN and instead headed ventrally, along the stria medullaris. We were able to follow one cell to the endopiriform cortex, but it did not form collaterals, suggesting the main axon continued to more posterior regions, which remain to be determined. We cannot rule out that tortuosa cells target retrohippocampal areas like other ADn cells but via this alternative route. Despite the axon reaching the cortex without passing or innervating the TRN, it is likely that these cells still receive feedback from GABAergic neurons of the anterodorsal TRN (Pinault & Deschenes, 1998; Vantomme *et al*., 2020), which may contribute to the burst firing patterns observed in tortuosa and other kinds of HD cells. Inactivating GABAergic presynaptic terminals of TRN neurons causes a net increase in ADn cell firing rates while preserving HD tuning (Duszkiewicz *et al*., 2024).

## Conclusions

Our data provide novel insights into the granularity of the mouse ADn in terms of HD cell firing patterns and connectivity. The advantage of our approach is that single HD cells could be targeted, extracellularly recorded, then juxtacellularly labeled, enabling us to determine the precise location within the ADn, the neurochemical profile (CR+ or CR-), and axon projection, linking these features to their firing patterns. Further recordings and labeling are required to determine whether e.g. ON excitation HD cells or sound-motion activated HD cells map on to specific types, or whether there is a degree of plasticity in the circuits. Nevertheless, the short-latency responses to light flashes and sounds demonstrate the ADn operates with separate parallel channels conveying HD-modulated stimuli to the rest of the HD network. We identified 3 types of projection pattern, but there are likely more if the main cortical target areas are included. Altogether, our results reveal a rich microarchitecture within the HD network. HD cells differ in molecular identity, connectivity, sensorimotor responsiveness, and firing characteristics, with CR+ cells forming a medio-lateral gradient that extends ventrally along the midline thalamus to reach the nucleus reuniens. Our findings suggest a re-evaluation of this HD hub as a network of parallel circuits, coordinating context-specific, sensory-modulated updates of the head direction for spatial navigation.

---

**Figure S1.**
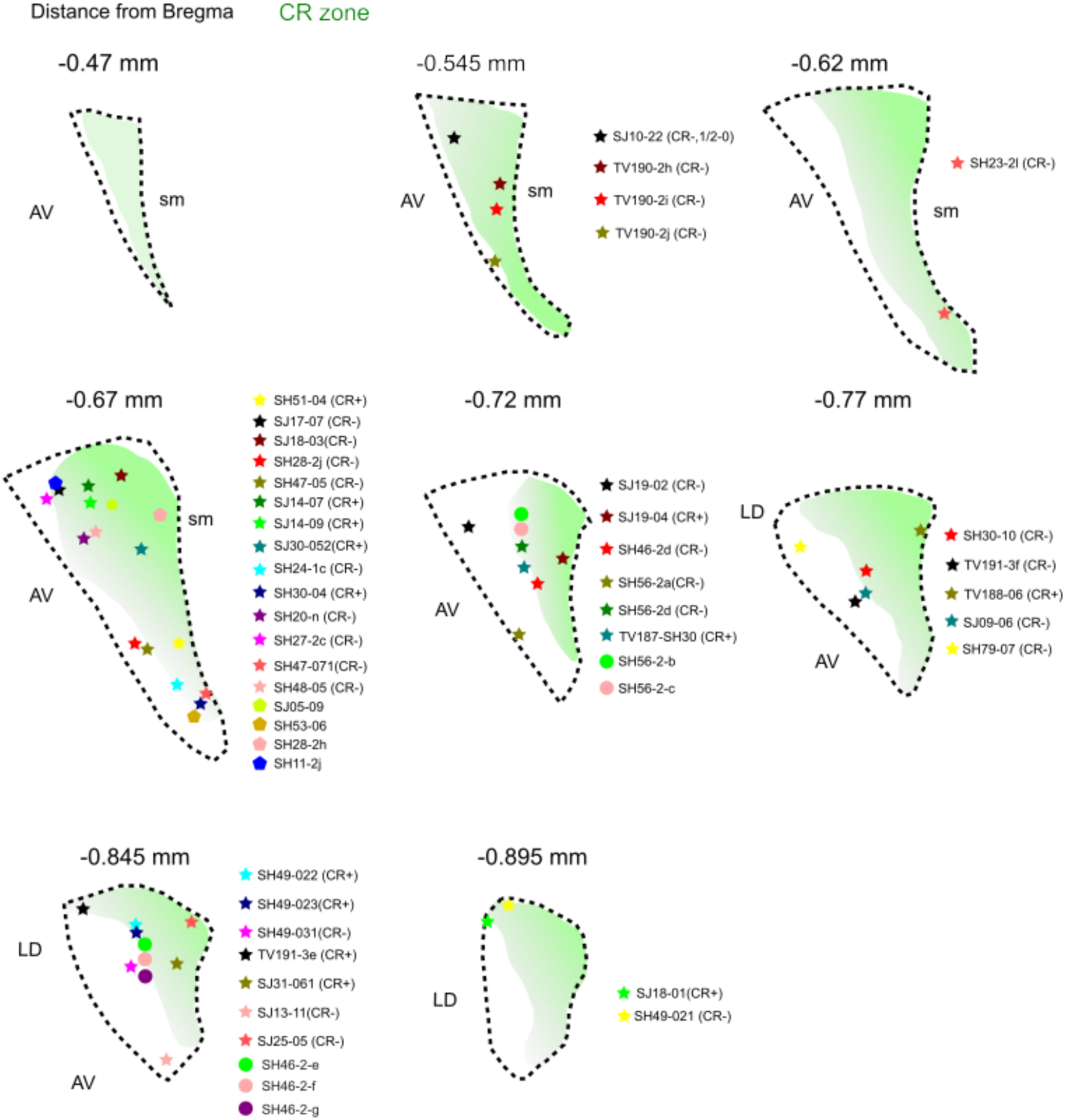
Recording locations of single extracellularly recorded ADn cells, related to Figure 1. Schematic map showing the locations of recorded ADn cells along with their names (stars, 40 labeled cells; pentagons, 4 partially labeled cells; circles, 6 unlabeled cells). Shaded areas indicate regions of CR immunoreactivity (‘CR zone’). Cells were named with the initials of the experimenter and the number of cells they recorded.

**Figure S2.**
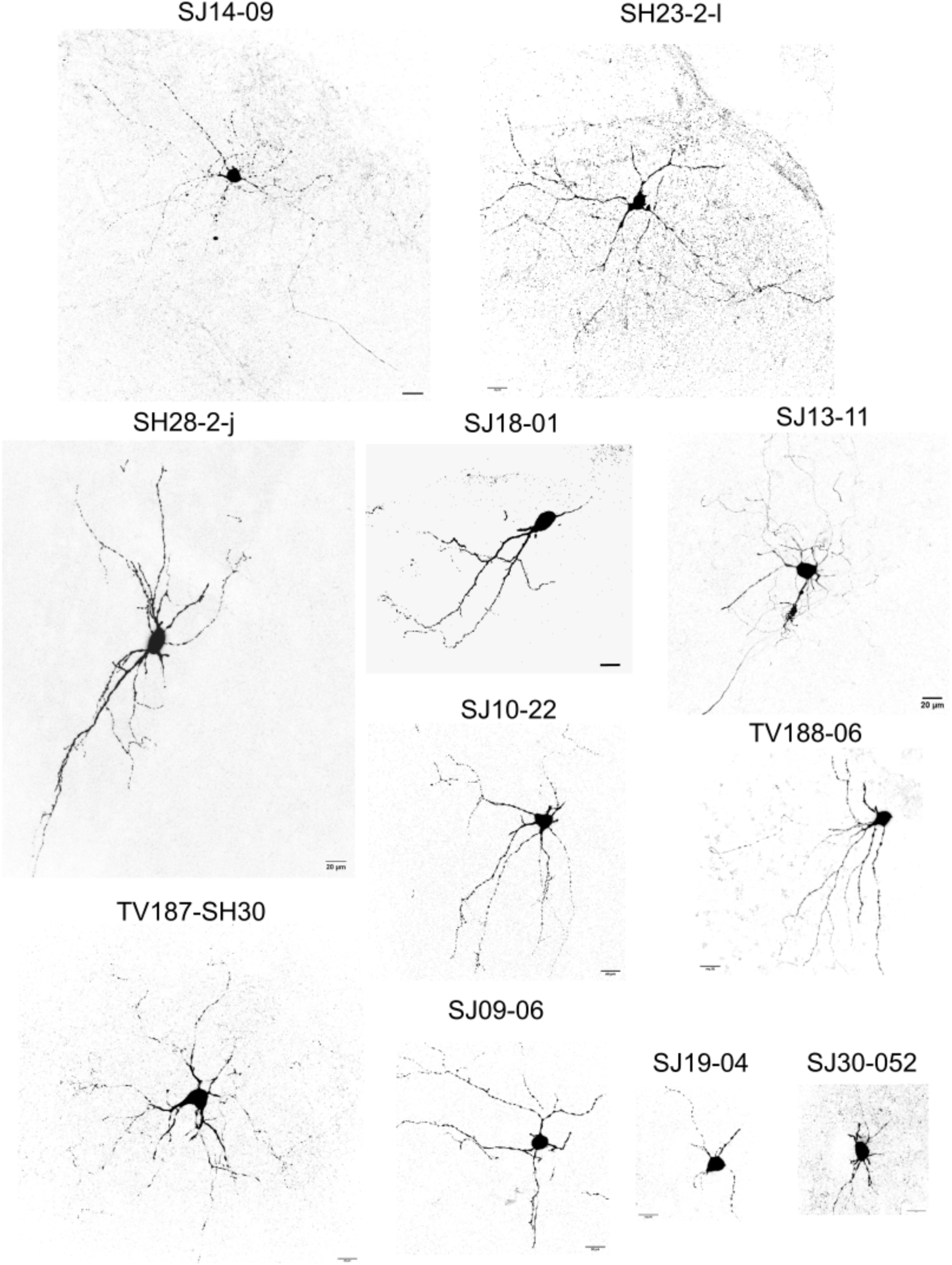
Images of juxtacellularly labeled ADn HD cells, related to Figure 1. Confocal maximum intensity z-projections of single 70 µm thick sections containing the soma and primary dendrites of labeled HD cells in the ADn. Scale bars, 20 µm. z-thickness (μm) for each cell: SH23-2l (47.5), SH27-2c (72), SH28-2-j (33.5), SJ14-09 (65.9), SJ18-01 (70), SJ10-22 (65.5), SJ13-11 (16.1), SJ09-06 (41.7), SJ19-04 (31.7), SJ30-052 (38), TV187-SH30 (66), TV188-06 (54.5).

**Figure S3.**
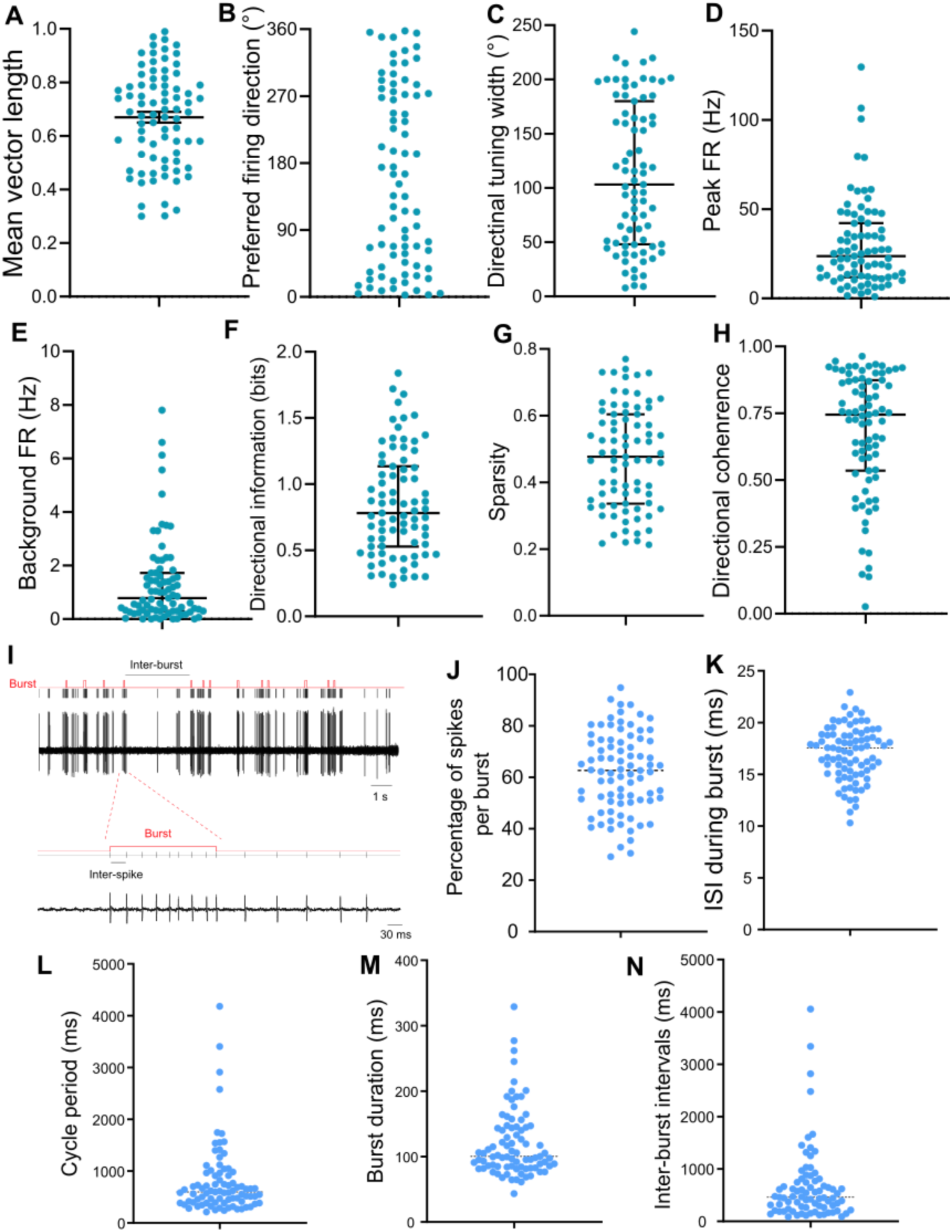
Basic firing characteristics and bursting patterns of recorded ADn HD cells, related to Figure 1. Properties of recorded HD cells. Left to right: mean vector length, PFD, directional firing range, peak firing rate (FR), background firing rate, directional information content, sparsity, and directional coherence. See Table S1 for further details of each cell. Data presented as median [IQR].

**Figure S4.**
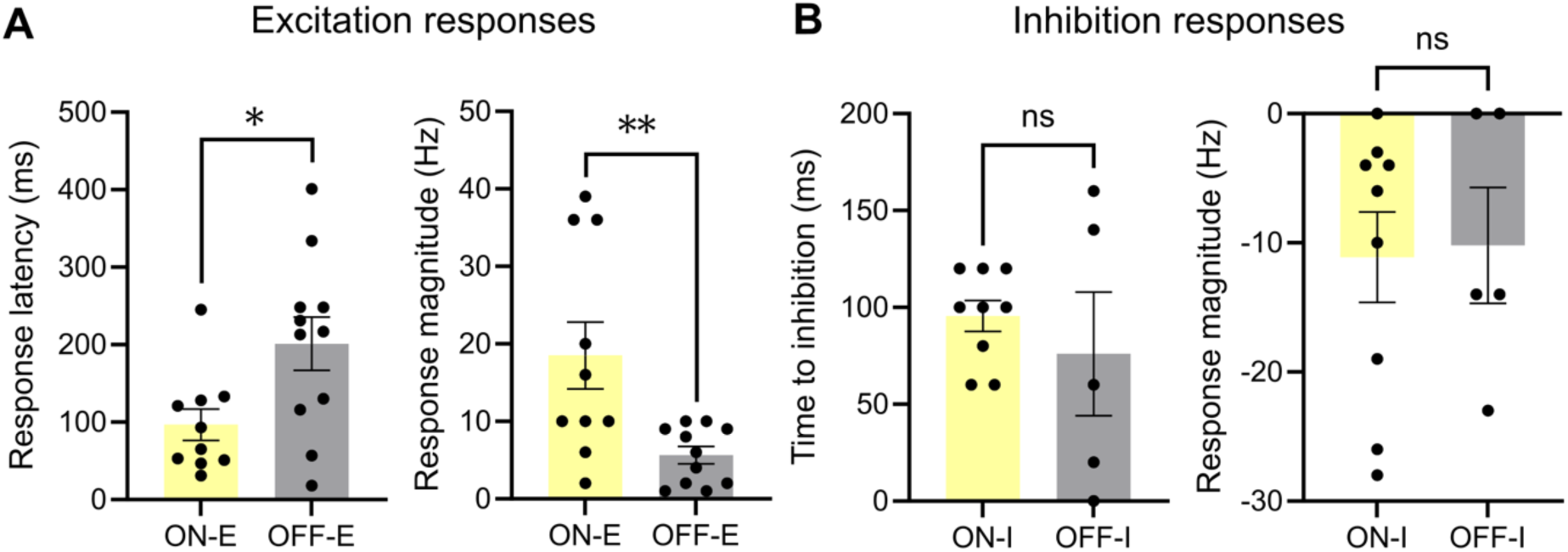
Comparison of ADn HD cell responses to light-ON and light-OFF, related to Figure 2. Response latency and magnitude after light-ON (yellow) or light-OFF (grey) for excitation responses (**A**) and inhibition responses (**B**), data presented with mean ± SEM, *p*\* < 0.05, *p*\*\* < 0.01, ns, not significantly different.

**Figure S5.**
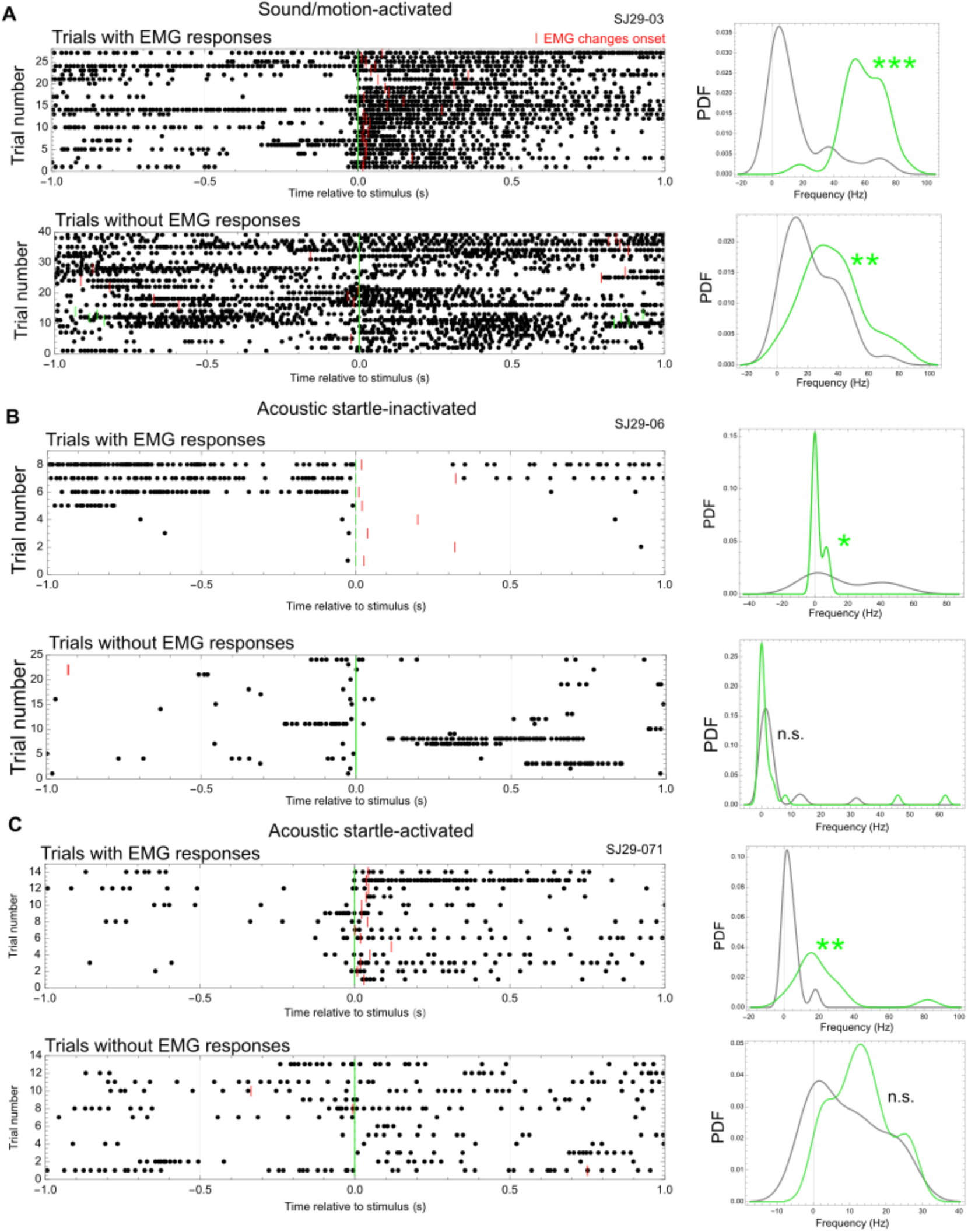
ADn HD cell responses to sensorimotor stimuli, related to Figure 3. Raster plots (left) and PDFs (right) showing responses to sound stimulus with and without EMG responses separately for cell SJ29-03 (**A**), SJ29-06 (**B**), and SJ29-071 (**C**). In raster plots, each point represents a spike, green ticks show other sound onset events, and red ticks mark the onset time of abrupt EMG increases.

**Figure S6.**
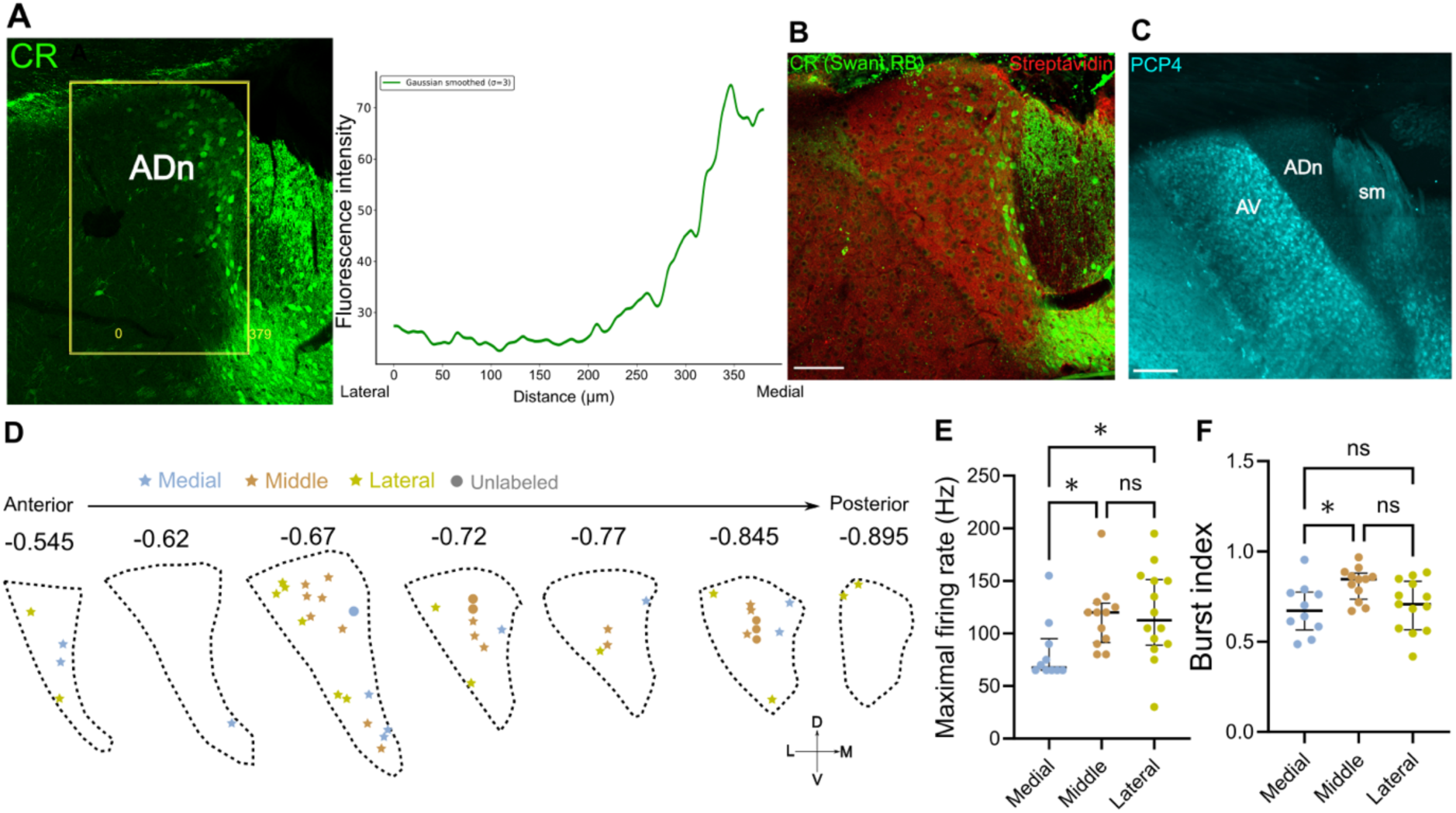
Distribution of calretinin immunoreactivity in the ADn, related to Figure 4. (**A**) Fluorescence intensity histogram for CR based on the ROI (boxed region). (**B**) Gradient in mouse ADn showed with another antibody for CR. (**C**) PCP4 immunoreactivity in mouse AV, case TV183. Scale bar, 200 μm. (**D**) Schematic map showing locations of identified ADn cells with groups they are divided into: Medial (blue), Middle (brown), Lateral (green). (**E**-**F**) Comparison of maximal firing rate (**E**) and burst index (**F**) between medial, middle, and lateral ADn HD cells.

**Figure S7.**
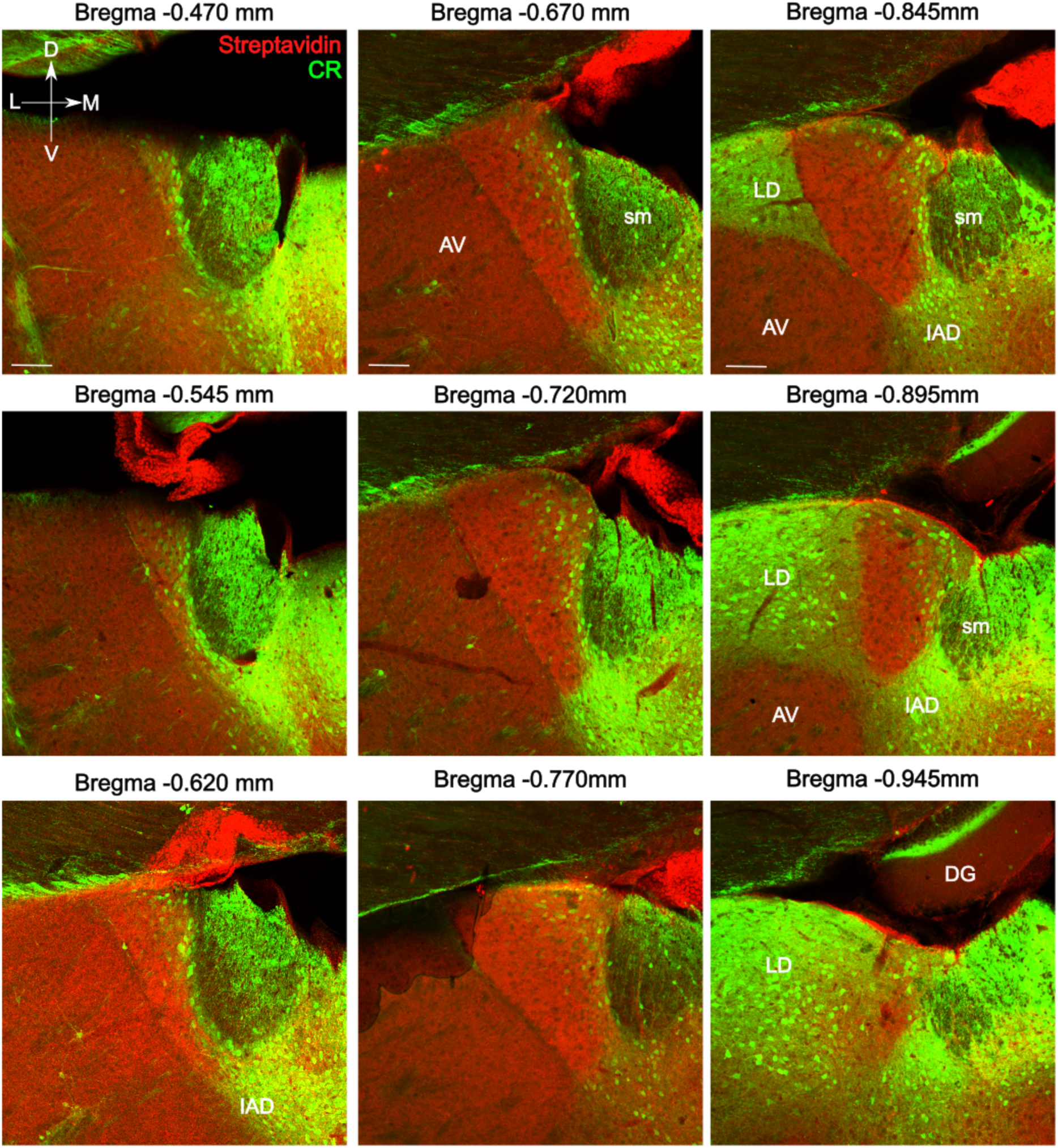
Serial mouse coronal sections tested for calretinin, related to Figure 4. Single optical confocal images. Sections from mouse TTPS8.5d, scale bars, 100 μm. L, lateral; M, medial; D, dorsal; V, ventral. AV, anteroventral thalamic nucleus; DG, dentate gyrus; LD, laterodorsal thalamic nucleus; IAD, interanterodorsal thalamus; sm, stria medullaris of the thalamus.

**Figure S8.**
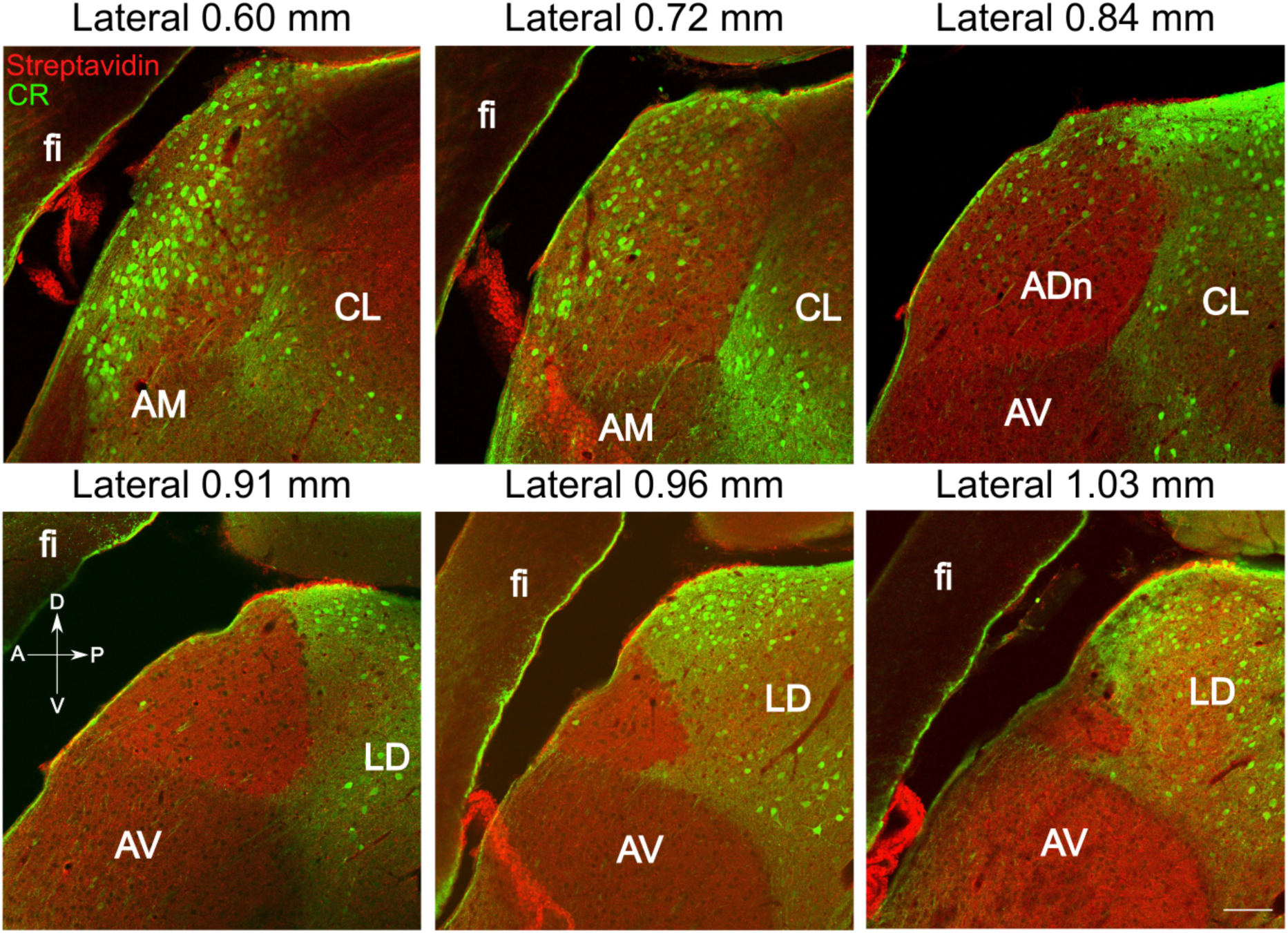
Serial mouse sagittal sections tested for calretinin, related to Figure 4. Single optical confocal images. Sections from mouse SJ33, scale bars, 100 μm. A, anterior; P, posterior; D, dorsal; V, ventral. AV, anteroventral thalamic nucleus; AM, anteromedial thalamic nucleus; CL, centrolateral thalamic nucleus; fi, fimbria.

**Figure S9.**
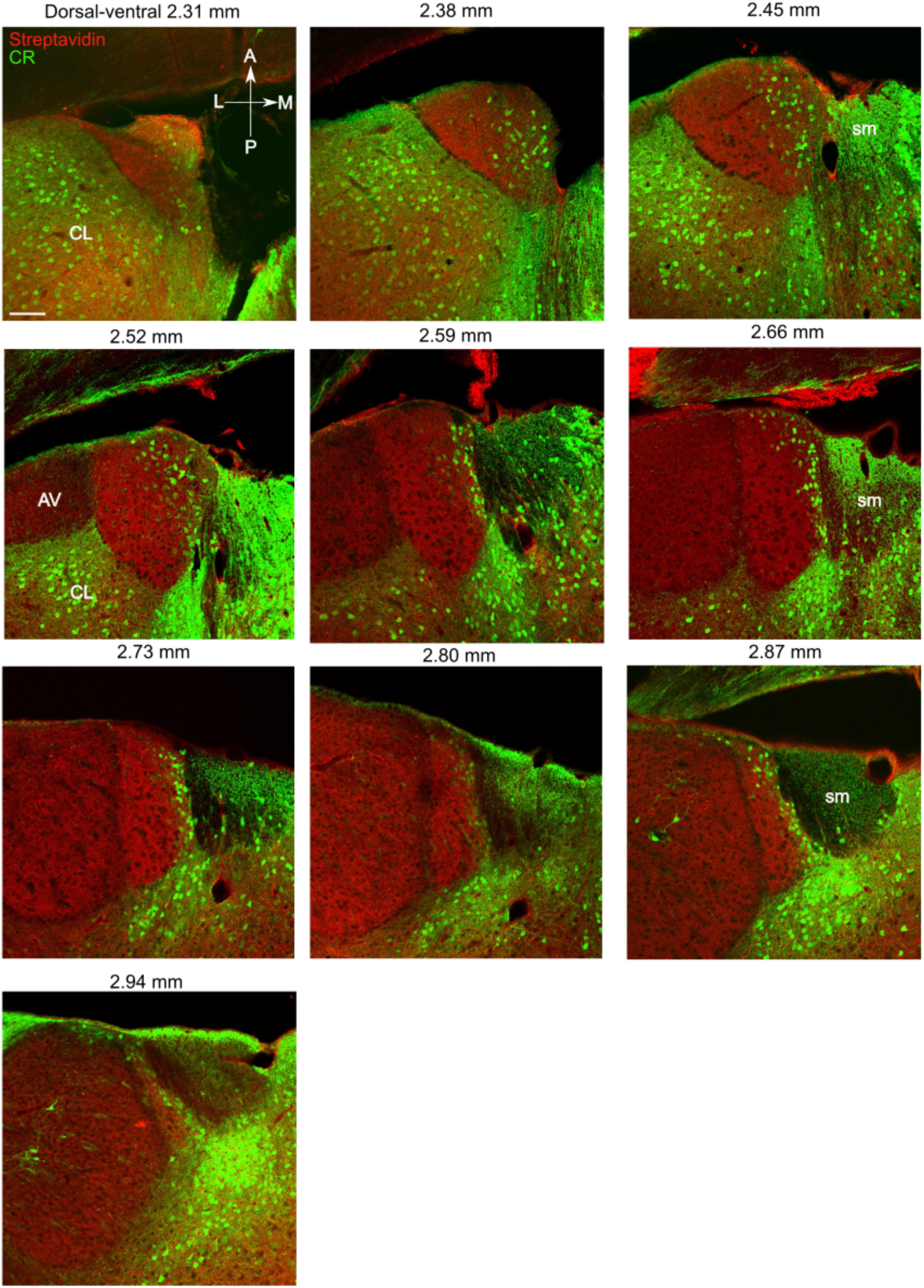
Serial mouse horizontal sections tested for calretinin, related to Figure 4. Single optical confocal images. Sections from mouse M311, scale bars, 100 μm. A, anterior; P, posterior; L, lateral; M, medial. AV, anteroventral thalamic nucleus; CL, centrolateral thalamic nucleus; sm, stria medullaris of the thalamus.

**Figure S10.**
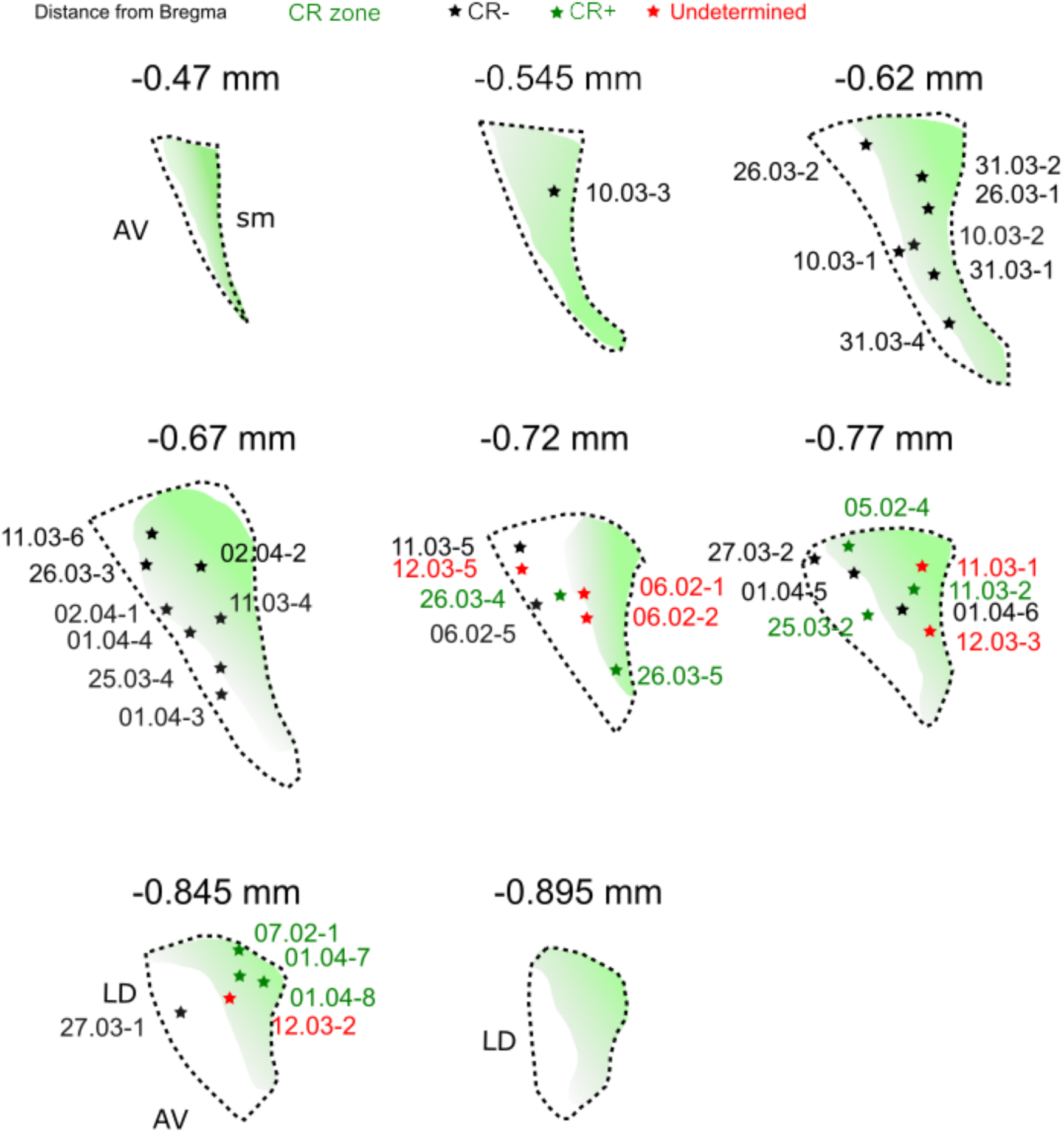
Map of ex vivo recorded ADn cells, related to Figure 4. Schematic map showing the locations of recorded ADn cells along with their names (green stars: labeled CR+ cells; black stars: labeled CR-cells; red stars: undetermined cells). Shaded areas indicate regions of CR immunoreactivity (‘CR zone’).

**Figure S11.**
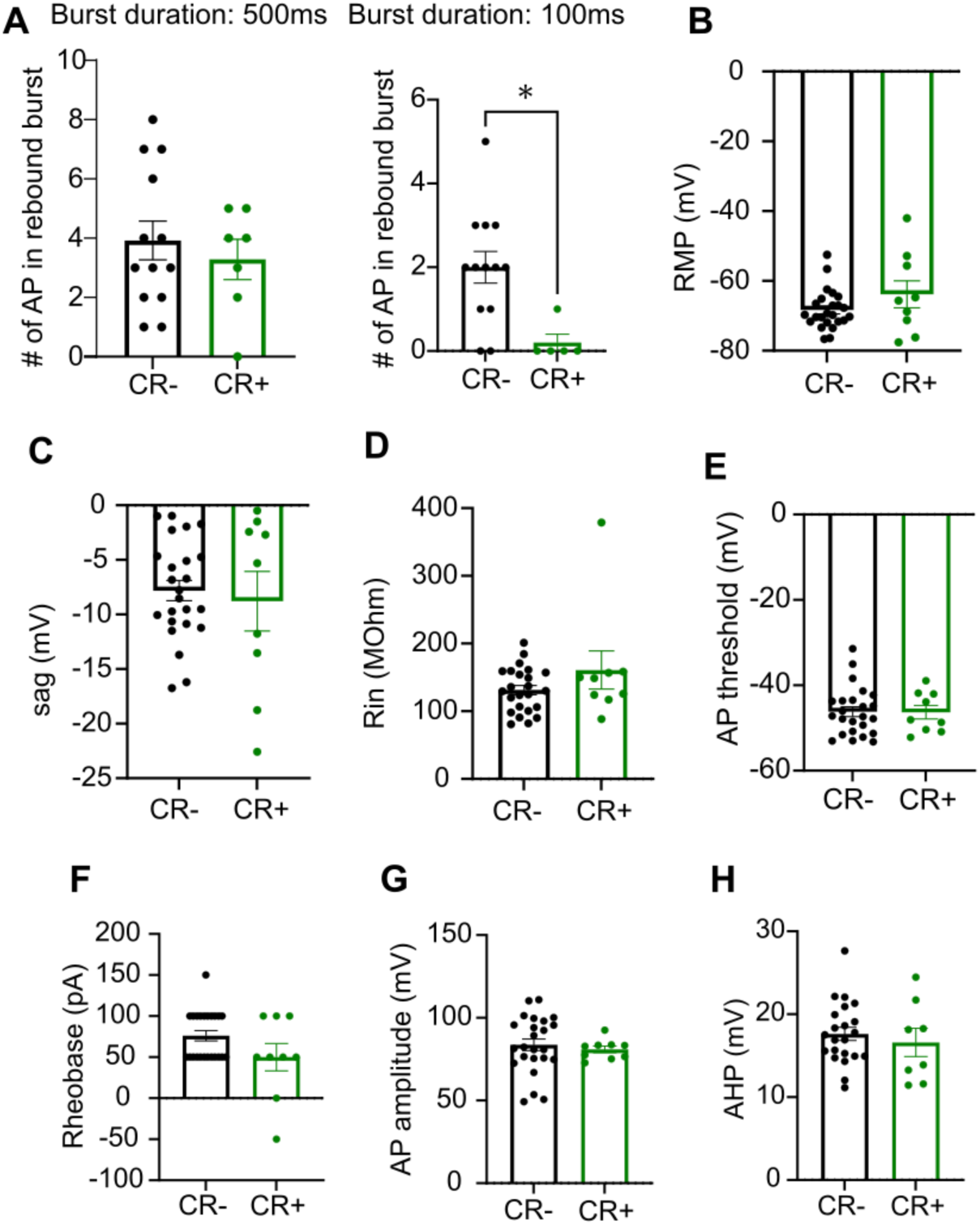
Intrinsic properties of CR+ and CR-ADn cells, related to Figure 4. (**A**) Number of action potential (AP) within the rebound burst following 500 ms (left) and 100 ms (right) hyperpolarizing current. (**B**) Resting membrane potential (RMP) in mV. (**C**)Sag ratio (mV). (**D**)Input Resistance (Rin) in MegaoOhm. (**E**) Action potential threshold in mV (AP). (**F**) Rheoabse in pA. (**G**) Action Potential amplitude in pA. (**H**) Afterhyperpolarization potential in mV. (n=13 CR- and n=7 CR+ cells from 8 mice).

## Supplementary Tables

**Table S1. Basic parameters of single recorded ADn HD cells, related to Figure 1.**

Firing properties of recorded HD cells along with their CR immunoreactivity. See Methods for analysis of each parameter. Cells listed with NA were confirmed as HD cells during recording but without available imu information. A total of 113 ADn cells (including 3 non-HD cells) were recorded from 35 mice.

**Table S2. Responses of ADn HD cells to light stimuli, related to Figure 2.**

A total of 27 ADn cells tested for responses to light stimuli are listed based on their response type. *Data presented as median, IQR. Abbreviations: IR, immunoreactivity; u, data unavailable; na, not applicable; WSR test, Wilcoxon signed-rank test.

**Table S3. Responses of ADn HD cells to sound stimuli, related to Figure 3**.

A total of 21 ADn cells tested for responses to sound stimuli at their PFDs or UPFDs are listed based on their response type. *Data presented as median, IQR. Abbreviations: IR, immunoreactivity; u, data unavailable; y, yes; n, no; na, not applicable; WSR test, Wilcoxon signed-rank test.

**Table S4. Comparisons of firing pattern and intrinsic properties CR+ and CR-ADn cells, related to Figure 4**.

Statistical reporting of basic firing properties (from *in vivo* recordings) and intrinsic properties (*ex vivo* recordings) of CR+ and CR-ADn cells. Mean, standard deviation (SD), SEM, and *p* values (unpaired t-tests or Mann-Whitney tests).

**Table S5. Features of labeled ADn cells, related to Figures 6-8.**

A total of 40 cells were labeled then recovered and localized to the ADn and examined for immunoreactivity to CR. A total of n=17 cells were recorded from wildtype (wt) mice, n=18 cells from mice injected with AAV-CBh-GFP, and n=5 from mice injected with an AAV encoding human mutant tau but lacked viral expression in the ADn. Three cells identified as glia cells were due to pipette damage to the cell after juxtacellular modulation. Only dendrites of cell SH28-2h were detected but not somata due to the missing sections. R/L, right/left hemisphere; labeling strength: - (unlabeled), + (only somata detected), ++ (somata and dendrites detected), +++ (somata, dendrites, and axon detected), ++++ (somata, dendrites, and axon detected in PrS); na, not available; u, unknown; aDMS, anterior DMS; cg, cingulum; ePir, endopiriform cortex. For projection patterns (types I, II, III), see Fig. 8H. Note, axon trajectory indicates some of the locations where the main axon was observed, irrespective of collaterals or terminals.

## Resource Availability

### Lead contact

Further information and requests for resources and reagents should be directed to and will be fulfilled by the lead contact, Tim Viney.

### Materials availability

Apart from the viral vectors (see Key resources table), this study did not generate new unique reagents.

### Data and code availability

Code will be made available on GitHub.

## Acknowledgements

We thank Barbara Sarkany, Aditi Athreya, and Kathryn Holland for help with tissue processing, Brook Perry for advice on implementing the IMU, and Martyn Preston for building the light module. We also thank Patvitra Reanchareonsuk for help with 2D reconstructions. Funding: Alzheimer’s Society grant 522 AS-PhD-19a-010 (T.J.V.); Medical Research Council grants MR/R011567/1 and MR/Z504518/1 (T.J.V.); The John Fell Fund grant 0013781 (T.J.V.); UKRI grant EP/Z001358/1 (T.J.V., S.H.); NIH R01 MH120073 (M.E.H.); U.S. Office of Naval Research MURI N00014-1-19-2571 (M.E.H.); NIH F32MH139270 (P.A.L.). S.H. was supported by a Blaschko Fellowship from the Department of Pharmacology, Oxford. S.J. was supported by a Clarendon Scholarship.

## Author Contributions

Conceptualization: S.H., S.J., T.J.V. Methodology: S.H., S.J., T.J.V. Software: S.H., S.J., P.A.L., T.J.V. Validation: S.H., S.J., P.A.L., M.E.H., T.J.V. Formal analysis: S.H., S.J., M.W., J.Q., T.J.V. Investigation: S.H., S.J., M.W., J.Q., P.A.L., M.E.H., T.J.V. Resources: S.H., M.E.H., T.J.V. Data curation: S.H., S.J., T.J.V. Writing—original draft: S.H., S.J., T.J.V. Writing—review and editing: S.H., S.J., M.W., J.Q., P.A.L., M.E.H., T.J.V. Visualization: S.H., S.J., M.W., J.Q., T.J.V. Supervision: T.J.V. Project administration: T.J.V. Funding acquisition: S.H., T.J.V.

## Methods

### Key resources table

| REAGENT or RESOURCE | SOURCE | IDENTIFIER |
| --- | --- | --- |
| **Antibodies** |  |  |
| CR | Swant | 7699/3H |
| CR | ABclonal | A21965 |
| PCP4 | Santa Cruz Biotech | G0814 |
| VGLUT1 | Synaptic System | 135304 |
| VGLUT2 | Synaptic Systems | Cat. No. 135 404, RRID:AB_887884 |
| Bassoon | Assay designs | VAM-PS003 |
| CHAT | Synaptic Systems | Cat.No. 297013 |
| CHAT | CHEMICON International | AB144P-200UL |
| Parvalbumin | Synaptic Systems | 195004/1-19, RRID: AB_2156476 |
| C1ql2 | Novus Biologicals | Cat. No.NBP2-34090 |
| NeuN | EMD Millipore Corp | Lot: 3713321 |
| **Chemicals, peptides, and recombinant proteins** |  |  |
| retroAAV-hSyn-EGFP | Addgene | Addgene: 50465 |
| RetroAAV-EF1a-DIO-EYFP | Addgene | Addgene: 27056 |
| Neurobiotin tracer | Vector Laboratories | #SP-1120; RRID: AB_2313575 |
| Isoflurane inhalation anesthesia | Teva Ltd. | https://products.tevauk.com/p/Category?id=185 |
| Vetergesic (0.3 mg/ml Buprenorphine) | Ceva<br>Marcain (Bupivacaine) | Aspen #PS09728 |
| Refobacin bone cement | Zimmer Biomet | #3003940002-3 |
| Tetric EvoFlow dental cement | Ivoclar Vivadent | A1 |
| Dil tracer | Life Technologies | #V22889 |
| Normal horse serum | Vector Laboratories | S-2000 |
| Normal goat serum | Vector Laboratories | S-1000 |
| **Critical commercial assays** |  |  |
| Vectastain ABC Elite kit | Vector Laboratories | Cat# PK6100; RRID: AB_2336819 |
| Vectashield Antifade Mounting Medium | Vector Laboratories | Cat# H-1000; RRID: AB_2336789 |
| **Experimental models: Organisms/strains** |  |  |
| C57BL6j mice | Charles River Laboratories | https://www.criver.com/ |
| C1QL2-IRES-Cre mice (C57BL/6-C1ql2<sup>em1(cre)Gfng/J</sup>) | Jackson Laboratory | https://www.jax.org/strain/036955 |
| **Software and algorithms** |  |  |
| FIJI (ImageJ) | NIH | https://imagej.net/software/fiji/ |
| MATLAB | Mathworks | https://www.mathworks.com/products/matlab.html |
| Python |  | https://www.python.org/ |
| Zen Black, Zen Blue, Axiovision | Zeiss | www.zeiss.co.uk |
| Spike2 | Cambridge Electronic Design | http://ced.co.uk/ |
| GraphPad Prism 7 | GraphPad Software | https://www.graphpad.com/ |
| Wolfram Mathematica |  | Wolfram Mathematica: Modern Technical Computing |

### Experimental model and subject details

All procedures involving experimental animals were under approved personal and project licenses according to UK Animals (Scientific Procedure) Act 1986 and associated regulations. Experiments were conducted with adult male and female C57Bl/6J mice and heterozygous C1QL2-IRES-Cre mice (C57BL/6-C1ql2^em1(cre)Gfng^/J, strain #036955, The Jackson Laboratory). A total of 59 mice were used in this study. We performed *in vivo* recordings from 45 mice, including 19 of these injected with AAVs reported in Jiang *et al*. (2024) (Table S1, S4). We performed *ex vivo* recordings in brain slices from 8 mice, performed additional tracing experiments in 3 mice, and conducted additional immunohistochemical tests in 3 mice.

#### Surgical procedures

*In vivo extracellular recordings in head-fixed mice*. Surgical procedures were performed as previously described (Viney *et al*., 2018; Viney *et al*., 2022). Four craniotomy sites and three screws sites were marked and drilled based on stereotaxic coordinates. Craniotomy sites (in mm from Bregma): hippocampal CA1, −2.5 to −2.3 antero-posterior (AP), +1.5 medio-lateral (ML), and −2.3 AP, −1.5 ML; ADn, −0.85 AP, ±0.75 ML. Two M1.2 x 3 screws (Precision Technology Supplies Ltd) with a soldered female pin were fixed to the skull above the cerebellum for ground/reference and securing the head plate. One or two M1.2 x 2 screws were fixed above the motor cortex (+1.5 AP, ±1.7 ML) to support the head plate. Screws were sealed with Refobacin bone cement (Zimmer Biomet), and a machined glass-reinforced plastic D-shaped headplate (0.7 g, custom made at the Department of Physics, Oxford University) was positioned over the screws and secured with the bone cement.

In the experiments for detecting startle responses evoked by a sudden sound, an electromyogram (EMG) was used to help detect movement of the animal from the neck muscle. A small incision was made in the skin at the dorsal neck region. A twisted stainless steel wire was positioned under the skin from the neck incision to the muscles.

*Anterograde and retrograde tracing.* ∼60 nl of AAVs (AAV-CBh>EGFP:WPRE, AAV8(VB900088-2238xse)-C, VectorBuilder; AAV1-hSyn1-DIO-EGFP-WPRE, VVF115) were bilaterally pressure-injected using glass pipettes into the ADn (−0.8 AP, 0.75 ML, 2.8 dorsoventral (DV) from brain surface),. For retrograde labeling of ADn cells: C57BL6j mice were injected with ∼400 nl of rAAV (RetroAAV-hSyn-EGFP, Addgene: 50465) at the following coordinates: −4.5 AP, 2.75 ML, 3.8 dorsoventral (DV) from brain surface. C1QL2-IRES-Cre mice were injected with ∼400 nl of rAAV (RetroAAV-Ef1a-DIO-EYFP, Addgene: 27056) at the following coordinates: −3 AP, 0.3 ML, 1.3 dorsoventral (DV) from brain surface. Following recovery, mice were kept for >1 week (AAV injections) until transcardial perfusion.

#### *In vivo* recordings and juxtacellular labeling

Glass electrode recordings in head-fixed mice were performed as previously described (Jiang *et al*., 2024) (Fig. 1). The main glass electrode (filled with 3% neurobiotin in 0.5 M NaCl) was lowered towards the ADn. Another glass electrode was lowered to the CA1 pyramidal cell layer. The experimenter manually rotated the setup to control the animal’s heading while dynamically adjusting the depth of the main glass electrode between 2 mm and 3.5 mm DV, in order to increase the chances of passing through the receptive fields of HD cells. The experimenter learned to recognize the specific sound (pitch, frequency, temporal pattern) of ADn HD cell spikes via a loudspeaker, along with the abrupt firing rate increase associated with entering a receptive field. Therefore, our dataset is biased towards HD cells versus non-HD cells in the ADn. Once an HD cell was detected, at least one full slow turn was conducted. If no HD cells were detected, the glass electrode was removed from the brain and inserted at a slightly different medio-lateral/antero-posterior position. This was conducted for both hemispheres during 2-4 h recording sessions over 1-4 days. The orientation of the mouse’s head was continuously acquired from an inertial measurement unit (IMU) datalogger (SparkFun OpenLog Artemis) attached to the apparatus, which was preprogrammed to automatically log data from the Global Navigation Satellite System.

Juxtacellular labeling was performed after completing recordings from all detected HD cells in a recording session. Neurobiotin was delivered via the glass electrode to a recorded cell of interest at the end of the recording session using 200 ms current pulses followed by a 4–7 h recovery period prior to perfusion.

#### Sensory stimulation

After detecting HD cells with stable recordings while the mouse was passively rotated, sensory stimuli were presented when the mouse was facing the PFD or was outside the PFD. The recording room was kept in photopic conditions (600-700 lux) when stimulation was delivered.

A customized LED strip (7 cm long) was attached to the setup facing the mouse’ head, 10.5 cm away from the animal, providing a viewing angle of approximate ∼30 deg on each side from the centre of the LED strip. The light stimulus consisted of light-ON (maximum luminance: 2,000-2,400 lux) and light-OFF (minimum luminance, i.e., room luminance: 150-200 lux), alternatingly presented for different durations (200 ms - 10 s) and trials (9 to 138 trials) per cell. This stimulus was used to assess transient or sustained responses. The ‘click’ sound (wide-broad frequency, intensity at mouse ear position: 45-73 dB, duration 10 ms) were recorded from a real finger click sound and played through a loudspeaker (fixed-position, 120 cm away from the mouse). The timing of light-ON/OFF for LED and sound stimuli were manually controlled with a customized module connected to the PC used for recordings. To deliver the LED or sound stimuli, the mouse was first positioned within or outside the PFD of the cell to compare their responses within and outside the receptive field.

#### Slice physiology

Whole-cell recordings of cells within ADn were performed as following: After decapitation, brains were placed in an ice-cold partial sucrose-based solution containing (in mM): sucrose 70, NaCl 70, NaHCO3 25, KCl 2.5, NaH2PO4 1.25, CaCl2 1, MgSO4 5, sodium ascorbate 1, sodium pyruvate 3, and D(+)-glucose 25 (carboxygenated with 5% CO2/95% O2; 305 mOsmol/kg). Coronal slices from the ADn (250 μm thick) were obtained with a vibrating slicer (Leica VT1200s). Next, the slices recovered at room temperature for 1 h incubated in holding artificial cerebrospinal fluid (ACSF) containing (in mM): 127 NaCl, 25 NaHCO3, 25 D(+)-glucose, 2.5 KCl, 1.25 NaH2PO4, 2 CaCl2, 3 sodium pyruvate, 1 sodium ascorbate, and 2 MgCl2 (carboxygenated with 5% CO2/95% O2; 310 mOsmol/kg). Slices were then transferred into the recording chamber where they were continuously perfused with recording ACSF (in mM): 127 NaCl, 25 NaHCO3, 25 D-glucose, 2.5 KCl, 1.25 NaH2PO4, 1 MgCl2, and 2 CaCl2 (310 mOsmol/kg). Cells were visualized using an upright microscope (BX51WI, Olympus) equipped with oblique illumination optics (WI-OBCD; numerical aperture 0.8) and a 40× water-immersion objective. Images were collected by a CCD camera (Oxford Instruments Andor Lt) operated by ImageJ software. ADn cells were identifed by their location and typical rebound-burst following a hyperpolarizing current. Electrophysiological recordings were acquired using HEKA EPC10 (10 Hz sampling rate) at 32°C with HEKA PATCHMASTER for data acquisition. Patch pipettes were pulled from borosilicate glass (Warner instruments) with an open tip of 3.5–5 MegaOhm of resistance and filled with intracellular solution containing (in mM) 125 K-gluconate, 10 NaCl, 2 Mg-ATP, 0.2 EGTA, 0.3 Na-GTP, 10 HEPES and 10 K2-phosphocreatine, pH 7.4, adjusted with KOH (280 mOsmol/kg), with 5 mg/mL biocytin (Sigma-Aldrich) to fill the cells. Series resistance was kept under 20 MOhm with correct bridge balance and capacitance fully compensated; cells that exceeded this value were not included in the study. Cells were filled with biocytin for at least 10 min.

Intrinsic passive and active membrane properties were recorded in current-clamp mode at resting membrane potential by injecting 500-ms of increasing current stimuli from −250 pA to +500 pA, at intervals of 50 pA. Data analysis was conducted using a custom-designed script in Igor Pro-9.0 (Wavemetrics).

Following the recordings, slices were fixed for 2 hours with 4% paraformaldehyde (PFA) and stored in 0.1 M phosphate buffer (PB; pH 7.4) at 4°C.

#### Histology

For tissue processing, procedures were performed as previously described (Viney *et al*., 2018; Viney *et al*., 2022).

*Transcranial perfusion and sectioning.* Mice were deeply anesthetized with sodium pentobarbital (50 mg/kg, i.p.) and transcardially perfused with saline followed by a fixative solution containing 4% PFA (w/v, Sigma-Aldrich), 15% saturated picric acid (v/v, Sigma-Aldrich), and 0.05 % glutaraldehyde (w/v, distilled grade, TAAB Laboratories Equipment Ltd) 0.1 M PB. Some brains were post-fixed overnight in fixative lacking glutaraldehyde. After washing out the fixative, brains were stored in 0.1 M PB with 0.05% sodium azide at 4°C. Brains were sectioned at 70 µm thickness using a vibrating microtome (VT 1000S vibratome, Leica Microsystems) and stored in 0.1 M PB with 0.05% sodium azide at 4°C.

For the visualization of neurobiotin-labeled processes with fluorescence microscopy, brain sections were permeabilized in Tris-buffered saline (0.9% NaCl buffered with 50 mM Tris, pH 7.4; TBS) with 0.3% Triton X-100 (TBS-Tx) or via rapid 2x freeze–thaw (FT) over liquid nitrogen. Cy3- or Cy5-conjugated streptavidin (Jackson ImmunoResearch) was applied at 1:500 dilution in TBS-Tx (or PB if permeabilized with FT) for 4 h at room temperature (RT) or overnight at 4°C. Sections were washed in TBS-Tx or TBS/PB (if permeabilized with FT) and mounted to glass slides in Vectashield (Vector Laboratories) and sealed with nail vanish. For the FT method, sections were first incubated for 4 hours in 20% (w/v) sucrose in 0.1 M PB.

Sections were transferred to a foil plate and were treated with two rounds of rapid freeze and thaw cycles over liquid nitrogen before being returned to the glass vials to be washed three times (each for 10 minutes) in 0.1 M PB.

For immunohistochemical tests, sections were initially blocked in TBS/TBS-Tx with 20% normal horse serum (NHS, Vector Laboratories) in TBS/TBS-Tx for 1 h. This was followed by incubating with primary antibodies with 1% NHS in TBS/TBS-Tx for 2-5 days at 4°C. Control sections were included that lacked the primary antibodies. Subsequently, sections were washed 3 times in TBS/TBS-Tx, then incubated with secondary antibodies in 1% NHS in TBS/TBS-Tx for 4 h RT or overnight at 4°C. The following secondary antibodies (and dilutions) were used in various combinations (all raised in donkey): anti-mouse Alexa Fluor 405 (1:250) from Invitrogen, anti-guinea pig DyLight 405 (706-475-148) (1:250), anti-guinea pig, anti-rabbit, and anti-mouse Alexa Fluor 647 (706-475-148, 711-605-152, 705-605-151) (1:500) from Jackson ImmunoResearch, and goat anti-rabbit Alexa Fluor 405 (1:250) from Invitrogen. After 3 washes (10 minutes per wash) in 0.1 M PB, sections were mounted to glass slides in Vectashield and sealed with nail vanish.

For diaminobenzidine (DAB)-based horseradish peroxidase (HRP) reactions to visualize neurobiotin-labeled processes with light microscopy, some sections were blocked for 10 min at RT in 1% hydrogen peroxide (H_2_O_2_) in 0.1 M PB. Next, sections were incubated overnight at 4°C in 1:100 biotinylated goat anti-rabbit IgG (BA-1000, Vector Laboratories) in TBS containing 1% NGS. After washing 3 times in TBS, sections were incubated for 3 days at 4°C in avidin-biotinylated HRP complex (Vectastain ABC Elite kit, Vector Laboratories) in TBS. Subsequently, peroxidase was visualized using a mix of 1% nickel ammonium sulphate, 0.4% ammonium-chloride, and 3,3-DAB (0.5 mg/ml, Sigma-Aldrich) developed with 0.01% H_2_O_2_. After washing in PB, sections were treated with 0.25% osmium tetroxide (OsO_4_, TAAB Laboratories Equipment Ltd, UK) in 0.1 M PB for 5 min. Next, after washing in 0.1 M PB at least four times, sections were transferred onto slides in chrome alum gelatin and dried in air. Sections were then incubated in fresh xylene for 10 min before being quickly mounted in DePeX mounting medium.

#### Microscopy

For documentation (tiles or z-stacks) and qualitative evaluation for brain regions of interest, an AXIO Observer Z1 microscope (LSM 710; Zeiss) equipped with Plan-Apochromat objectives (magnification/numerical aperture: 10x/0.3, 20x/0.8 and 40x/1.4) was used to acquire images (Axiovision or ZEN Blue 2.6 software) across different fluorescence channels. For confocal microscopy, DIC M27 Plan-Apochromat 20x/0.8, 40x/1.4, 63×/1.4 and alpha Plan-Apochromat 100×/1.46 objectives were used. The following channel specifications were used for the detection of Alexa405, Alexa488/EYFP, Cy3, and Cy5: 405 nm: 405-30 solid-state laser, attenuation filter ND04, MBS-405, emission spectral filter 409–499 nm; 488 nm: Argon laser, MBS-488, emission spectral filter 493–542 nm; 543 nm: HeNe laser, MBS-458/543, emission spectral filter 552–639 nm; 633 nm: HeNe laser, MBS-488/543/633, emission spectral filter 637–757 nm. The pinhole was set to ∼1 Airy Unit for each channel to maintain a consistent optical slice thickness (0.6-0.7 μm) across all channels. Channels were acquired sequentially with beamsplitters set to minimize spectral overlap between channels (ZEN Black 14.0 software).

Neurons were traced using a drawing tube attached to a light microscope (Leitz Dialux22, Leica). After alignment of the tracings of consecutive sections, drawings were overlaid and copied onto a single sheet of tracing paper then digitized.

#### Quantification of ADn cells

ADn cells were imaged using tiled z-stacks that captured the entire ADn, with a step size of 1 µm, using a 40x/1.3 NA (oil immersion) objective. Colocalization of GFP+ cells, CR+ cells and DAPI in the ADn was quantified using the Cell Counter plugin in ImageJ based on 5 sections per mouse from n=3 mice. The proportion of CR+ cells was calculated by dividing the number of CR+ cells by the total number of DAPI+ nuclei. The small glial cell nuclei were excluded using a size threshold.

#### Analysis of electrophysiological data

*Head direction classification.* Neurons that exhibited bursting non-rhythmic firing that was clearly related to the animal’s head direction were initially selected for recording, which biased the initial search strategy to HD cells of the ADn. By visualizing labeled cells, we confirmed that recorded but unlabeled cells from either the same penetration site or from closely aligned coordinates were located in the ADn. Spikes were isolated by thresholding high-pass filtered voltage traces of their peaks and validated by using principal component analysis and/or visual inspection in Spike2 software (Cambridge Electronic Design, Cambridge, UK). In total, we recorded 143 ADn cells. We analyzed recordings that were longer than 100 s and with clear spikes isolated. A total of 26 cells were excluded because of lacking full coverage of 360° angles, and 21 cells were excluded due to unavailable IMU data or unclear spikes.

The directional tuning curve for each cell was obtained by plotting the firing rate as a function of the mouse’s directional heading, divided into bins of 6°. The firing rate was computed based on the total number of spikes divided by the total time in that bin. All HD properties-related parameters were analyzed based on published parameters (Clark *et al*., 2012) using custom Python scripts. From the directional tuning curve, we computed four parameters: 1) preferred firing direction. A Gaussian function (Blair *et al*., 1997) was used to approximate the HD at which the highest firing rate occurred to avoid deviations caused by small fluctuations in the raw directional tuning curve; 2) peak firing rate. The highest firing rate of the directional tuning curve rate, which indicates the firing rate when the mouse is facing in the cell’s preferred direction; 3) directional tuning width. The range of head directions over which the cell fires, which is defined as 6 times of the standard deviation to match the best-fit triangular function method (Taube *et al*., 1990); and 4) background firing rate. The average firing rate when the mouse is facing outside of the directional firing range of the cell.

The mean vector length (Rayleigh’s *r*) is a measure of the non-uniformity (or directionality) of the directional tuning curve and can vary between 0 (a uniform distribution) and 1 (a non-uniform distribution). Based on published criterion and subjective assessment of the directional tuning curve, a criterion of *r* > 0.30 was determined to HD cell. Other parameters analyzed include (1) Directional coherence. Directional coherence is a measure of the smoothness in the firing rate versus HD tuning curve. The firing rate for each directional bin is correlated to the firing rates in the two immediately adjacent neighbouring bins (CW and CCW directions) and an overall correlation is calculated across all bins. This measure would be high for cells that have a strong, continuous, and smooth looking tuning curve and low for cells that have a jagged, irregular, and uneven looking tuning curve. (2) Directional information content.

Directional information content is a measure of how many bits of HD information is conveyed by each spike and was calculated by the following formula: directional information content = p_i_ (l_i_/l) log2 (l_i_/l), where p_i_ is the probability that the head pointed in the i_th_ directional bin, li is the mean firing rate for bin i, and l is the mean firing rate across all directional bins. (3) Burst index. To measure the extent of burst spiking by HD cells, a burst index score was calculated for each cell. I first produced ISI histograms (0–100 ms) from the spike timestamped data for each cell. Next, I computed a burst index score by counting the number of ISIs that were <10 ms, but >2 ms due to the refractory period, and then dividing this value by the total number of ISIs that occurred between 0 and 100 ms. For this calculation I only used spikes that occurred when the animal’s head was within ±30° of the cell’s PFD (60° range). Thus, an HD cell that discharges in bursts would have a burst index measure closer to 1. (4) Coefficient of variation. CV is used to investigate whether there were differences in the regularity of spiking across HD cells from each brain area. CV is calculated by dividing the standard deviation of all the ISIs by the mean ISI.

Another type of plot showing number of spikes versus time was constructed to compute two parameters: 1) maximum firing rate, which is defined as the highest firing rate based on the 200 ms sample of the whole recording session; 2) mean firing rate, which is the average firing rate of the cell over the entire recording session.

*Responses to stimuli*. To investigate responses to light flashes and sound, we analyzed 0.5 s time windows before and after each stimulus using custom code in Mathematica (Wolfram Research Inc.). We report the following parameters (Tables S2 and S3): response frequency, median firing rate after the stimulus; Wilcoxon signed-rank test, comparing the number of spikes before versus after the stimulus (with an alpha value of 0.05); response magnitude, the difference between the median firing rates before and after the stimulus; response latency, median time to the first spike after the stimulus. Time to inhibition was represented as the median time differences between the stimulus onset (LED-ON/OFF and sound-ON) and the dropping time of the firing rate (when it was lower than Firing rate [Pre], bin size = 20 ms). Inclusion criteria: more than at least 6 presented stimuli per cell. In some cases, we separately analyzed firing patterns within the PFD and outside the PFD (UPFD). We also analyzed EMG in relation to sound stimuli using a similar approach.

### Statistics

All data are represented as mean ± SEM or median [IQR]. Experimental units (e.g. mice, cells) are specified in the text after the n values. Statistical analysis was carried out in GraphPad Prism, Python, IgorPro and Mathematica. The alpha was set to 0.05. For data that approximated a normal distribution (tested by the Shapiro-Wilk test), unpaired Student’s t tests were used to compare two groups with equal variances and unpaired t tests with Welch’s correction were used in two groups having different variances, otherwise Mann-Whitney tests were used. For non-normal distribution data, Wilcoxon signed-rank test was used to assess whether the population mean ranks differ. For comparisons of more than two groups we used Analysis of Variance (ANOVA) for parametric data followed by Tukey’s *post-hoc* test, and Kruskal-Wallis test for non-parametric data followed by Dunn’s test.

## References

Alexander, A.S., Robinson, J.C., Stern, C.E. & Hasselmo, M.E. (2023) Gated transformations from egocentric to allocentric reference frames involving retrosplenial cortex, entorhinal cortex, and hippocampus. Hippocampus, 33, 465–487.

Blair, H.T., Lipscomb, B.W. & Sharp, P.E. (1997) Anticipatory time intervals of head-direction cells in the anterior thalamus of the rat: implications for path integration in the head-direction circuit. J Neurophysiol, 78, 145–159.

Blanco-Hernandez, E., Balsamo, G., Preston-Ferrer, P. & Burgalossi, A. (2024) Sensory and behavioral modulation of thalamic head-direction cells. Nat Neurosci, 27, 28–33.

Brandon, M.P., Bogaard, A.R., Schultheiss, N.W. & Hasselmo, M.E. (2013) Segregation of cortical head direction cell assemblies on alternating theta cycles. Nat Neurosci, 16, 739–748.

Buzsaki, G. & Moser, E.I. (2013) Memory, navigation and theta rhythm in the hippocampal-entorhinal system. Nat Neurosci, 16, 130–138.

Clark, B.J., Harris, M.J. & Taube, J.S. (2012) Control of anterodorsal thalamic head direction cells by environmental boundaries: comparison with conflicting distal landmarks. Hippocampus, 22, 172–187.

Clark, B.J. & Harvey, R.E. (2016) Do the anterior and lateral thalamic nuclei make distinct contributions to spatial representation and memory? Neurobiology of Learning and Memory, 133, 69–78.

Clark, B.J., LaChance, P.A., Winter, S.S., Mehlman, M.L., Butler, W., LaCour, A. & Taube, J.S. (2024) Comparison of head direction cell firing characteristics across thalamo-parahippocampal circuitry. Hippocampus, **n/a**.

Clasca, F. (2023) Thalamic Output Pathways. In Clasca, F., Hofer, S.B., Usrey, W.M., Sherman, S.M. (eds) The Cerebral Cortex and Thalamus. Oxford University Press, pp. 121–131.

Conrad, C.D. & Stumpf, W.E. (1975) Direct visual input to the limbic system: Crossed retinal projections to the nucleus anterodorsalis thalami in the tree shrew. Experimental Brain Research, 23, 141–149.

Duszkiewicz, A.J., Orhan, P., Skromne Carrasco, S., Brown, E.H., Owczarek, E., Vite, G.R., Wood, E.R. & Peyrache, A. (2024) Local origin of excitatory–inhibitory tuning equivalence in a cortical network. Nature Neuroscience.

Farrow, K., Teixeira, M., Szikra, T., Viney, T.J., Balint, K., Yonehara, K. & Roska, B. (2013) Ambient illumination toggles a neuronal circuit switch in the retina and visual perception at cone threshold. Neuron, 78, 325–338.

Fernandez, D.C., Fogerson, P.M., Lazzerini Ospri, L., Thomsen, M.B., Layne, R.M., Severin, D., Zhan, J., Singer, J.H., Kirkwood, A., Zhao, H., Berson, D.M. & Hattar, S. (2018) Light Affects Mood and Learning through Distinct Retina-Brain Pathways. Cell, 175, 71–84.e18.

Gibson, B., Butler, William N. & Taube, Jeffery S. (2013) The Head-Direction Signal Is Critical for Navigation Requiring a Cognitive Map but Not for Learning a Spatial Habit. Current Biology, 23, 1536–1540.

Gong, J., Jellali, A., Mutterer, J., Sahel, J.A., Rendon, A. & Picaud, S. (2006) Distribution of vesicular glutamate transporters in rat and human retina. Brain Res, 1082, 73–85.

Grieves, R.M., Shinder, M.E., Rosow, L.K., Kenna, M.S. & Taube, J.S. (2022) The Neural Correlates of Spatial Disorientation in Head Direction Cells. eneuro, 9, ENEURO.0174-0122.2022.

Guillery, R.W. (1956) Degeneration in the post-commissural fornix and the mamillary peduncle of the rat. J Anat, 90, 350–370.

Hayakawa, T. & Zyo, K. (1989) Retrograde double-labeling study of the mammillothalamic and the mammillotegmental projections in the rat. J Comp Neurol, 284, 1–11.

Hinman, J.R., Brandon, M.P., Climer, J.R., Chapman, G.W. & Hasselmo, M.E. (2016) Multiple Running Speed Signals in Medial Entorhinal Cortex. Neuron, 91, 666–679.

Hinman, J.R., Chapman, G.W. & Hasselmo, M.E. (2019) Neuronal representation of environmental boundaries in egocentric coordinates. Nature Communications, 10, 2772.

Hintiryan, H., Rudd, M., Nanda, S., Gutierrez, A.E., Lo, D., Boesen, T., Garcia, L., Sun, J., Estrada, C., Mun, H.S., Yamashita, S., Han, Y.E., Bowman, I., Gou, L., Cao, C., Gonzalez, J., Moradi, K., Zhao, Q., Yenokian, I., Dev, A., Zingg, B., Xu, H., Xue, Q., Zhu, M., Liu, L., Chen, X., Yun, Z., Peng, H., Foster, N.N. & Dong, H.W. (2025) Distinct subnetworks of the mouse anterior thalamic nuclei. Nat Commun, 16, 6018.

Jankowski, M.M., Passecker, J., Islam, M.N., Vann, S., Erichsen, J.T., Aggleton, J.P. & O’Mara, S.M. (2015) Evidence for spatially-responsive neurons in the rostral thalamus. Front Behav Neurosci, 9, 256.

Ji, Z., Lomi, E., Jeffery, K., Mitchell, A.S. & Burgess, N. (2025) Phase Precession Relative to Turning Angle in Theta-Modulated Head Direction Cells. Hippocampus, 35, e70008.

Jiang, S., Hijazi, S., Sarkany, B., Gautsch, V.G., LaChance, P.A., Hasselmo, M.E., Bannerman, D. & Viney, T.J. (2024) Pathological tau alters head direction signaling and induces spatial disorientation. bioRxiv, 2024.2011.2007.622548.

Kapustina, M., Zhang, A.A., Tsai, J.Y.J., Bristow, B.N., Kraus, L., Sullivan, K.E., Erwin, S.R., Wang, L., Stach, T.R., Clements, J., Lemire, A.L. & Cembrowski, M.S. (2024) The cell-type-specific spatial organization of the anterior thalamic nuclei of the mouse brain. Cell Rep, 43, 113842.

Knierim, J.J., Kudrimoti, H.S. & McNaughton, B.L. (1995) Place cells, head direction cells, and the learning of landmark stability. J Neurosci, 15, 1648–1659.

Kropff, E., Carmichael, J.E., Moser, M.B. & Moser, E.I. (2015) Speed cells in the medial entorhinal cortex. Nature, 523, 419–424.

Lara-Vásquez, A., Espinosa, N., Durán, E., Stockle, M. & Fuentealba, P. (2016) Midline thalamic neurons are differentially engaged during hippocampus network oscillations. Scientific reports, 6, 29807–29807.

Lomi, E., Jeffery, K.J. & Mitchell, A.S. (2023) Convergence of location, direction, and theta in the rat anteroventral thalamic nucleus. iScience, 26, 106993.

Matyas, F., Komlosi, G., Babiczky, A., Kocsis, K., Bartho, P., Barsy, B., David, C., Kanti, V., Porrero, C., Magyar, A., Szucs, I., Clasca, F. & Acsady, L. (2018) A highly collateralized thalamic cell type with arousal-predicting activity serves as a key hub for graded state transitions in the forebrain. Nat Neurosci, 21, 1551–1562.

Mehlman, M.L., Winter, S.S. & Taube, J.S. (2019a) Functional and anatomical relationships between the medial precentral cortex, dorsal striatum, and head direction cell circuitry. II. Neuroanatomical studies. J Neurophysiol, 121, 371–395.

Mehlman, M.L., Winter, S.S., Valerio, S. & Taube, J.S. (2019b) Functional and anatomical relationships between the medial precentral cortex, dorsal striatum, and head direction cell circuitry. I. Recording studies. J Neurophysiol, 121, 350–370.

Mimura, Y., Mogi, K., Kawano, M., Fukui, Y., Takeda, J., Nogami, H. & Hisano, S. (2002) Differential expression of two distinct vesicular glutamate transporters in the rat retina. Neuroreport, 13, 1925–1928.

Morin, L.P. & Studholme, K.M. (2014) Retinofugal projections in the mouse. The Journal of comparative neurology, 522, 3733–3753.

O’Keefe, J. & Nadel, L. (1978) The Hippocampus as a Cognitive Map. Oxford: Clarendon Press.

Peyrache, A., Duszkiewicz, A.J., Viejo, G. & Angeles-Duran, S. (2019) Thalamocortical processing of the head-direction sense. Progress in Neurobiology, 183, 101693.

Pinault, D. & Deschenes, M. (1998) Projection and innervation patterns of individual thalamic reticular axons in the thalamus of the adult rat: a three-dimensional, graphic, and morphometric analysis. J Comp Neurol, 391, 180–203.

Piscopo, D.M., El-Danaf, R.N., Huberman, A.D. & Niell, C.M. (2013) Diverse visual features encoded in mouse lateral geniculate nucleus. J Neurosci, 33, 4642–4656.

Roska, B. & Werblin, F. (2001) Vertical interactions across ten parallel, stacked representations in the mammalian retina. Nature, 410, 583–587.

Sárkány, B., Dávid, C., Hortobágyi, T., Gombás, P., Somogyi, P., Acsády, L. & Viney, T.J. (2024) Early and selective localization of tau filaments to glutamatergic subcellular domains within the human anterodorsal thalamus. Acta Neuropathologica, 147, 98.

Shibata, H. (1993a) Direct projections from the anterior thalamic nuclei to the retrohippocampal region in the rat. J Comp Neurol, 337, 431–445.

Shibata, H. (1993b) Efferent projections from the anterior thalamic nuclei to the cingulate cortex in the rat. J Comp Neurol, 330, 533–542.

Sripanidkulchai, K. & Wyss, J.M. (1986) Thalamic projections to retrosplenial cortex in the rat. Journal of Comparative Neurology, 254, 143–165.

Stackman, R.W. & Taube, J.S. (1998) Firing Properties of Rat Lateral Mammillary Single Units: Head Direction, Head Pitch, and Angular Head Velocity. The Journal of Neuroscience, 18, 9020–9037.

Taube, J.S. (1995) Head direction cells recorded in the anterior thalamic nuclei of freely moving rats. J Neurosci, 15, 70–86.

Taube, J.S. (2007) The head direction signal: origins and sensory-motor integration. Annu Rev Neurosci, 30, 181–207.

Taube, J.S., Muller, R.U. & Ranck, J.B., Jr. (1990) Head-direction cells recorded from the postsubiculum in freely moving rats. I. Description and quantitative analysis. J Neurosci, 10, 420–435.

Tsanov, M., Chah, E., Vann, S.D., Reilly, R.B., Erichsen, J.T., Aggleton, J.P. & O’Mara, S.M. (2011) Theta-modulated head direction cells in the rat anterior thalamus. J Neurosci, 31, 9489–9502.

Tukker, J.J., Tang, Q., Burgalossi, A. & Brecht, M. (2015) Head-Directional Tuning and Theta Modulation of Anatomically Identified Neurons in the Presubiculum. J Neurosci, 35, 15391–15395.

Vann, S.D., Saunders, R.C. & Aggleton, J.P. (2007) Distinct, parallel pathways link the medial mammillary bodies to the anterior thalamus in macaque monkeys. Eur J Neurosci, 26, 1575–1586.

Vantomme, G., Rovó, Z., Cardis, R., Béard, E., Katsioudi, G., Guadagno, A., Perrenoud, V., Fernandez, L.M.J. & Lüthi, A. (2020) A Thalamic Reticular Circuit for Head Direction Cell Tuning and Spatial Navigation. Cell Rep, 31, 107747.

Viena, T.D., Rasch, G.E., Silva, D. & Allen, T.A. (2021) Calretinin and calbindin architecture of the midline thalamus associated with prefrontal-hippocampal circuitry. Hippocampus, 31, 770–789.

Viney, T.J., Salib, M., Joshi, A., Unal, G., Berry, N. & Somogyi, P. (2018) Shared rhythmic subcortical GABAergic input to the entorhinal cortex and presubiculum. Elife, 7.

Viney, T.J., Sarkany, B., Ozdemir, A.T., Hartwich, K., Schweimer, J., Bannerman, D. & Somogyi, P. (2022) Spread of pathological human Tau from neurons to oligodendrocytes and loss of high-firing pyramidal neurons in aging mice. Cell Rep, 41, 111646.

Vollan, A.Z., Gardner, R.J., Moser, M.-B. & Moser, E.I. (2025) Left–right-alternating theta sweeps in entorhinal–hippocampal maps of space. Nature.

Winnubst, J., Bas, E., Ferreira, T.A., Wu, Z., Economo, M.N., Edson, P., Arthur, B.J., Bruns, C., Rokicki, K., Schauder, D., Olbris, D.J., Murphy, S.D., Ackerman, D.G., Arshadi, C., Baldwin, P., Blake, R., Elsayed, A., Hasan, M., Ramirez, D., Dos Santos, B., Weldon, M., Zafar, A., Dudman, J.T., Gerfen, C.R., Hantman, A.W., Korff, W., Sternson, S.M., Spruston, N., Svoboda, K. & Chandrashekar, J. (2019) Reconstruction of 1,000 Projection Neurons Reveals New Cell Types and Organization of Long-Range Connectivity in the Mouse Brain. Cell, 179, 268–281.e213.

Yeomans, J.S., Li, L., Scott, B.W. & Frankland, P.W. (2002) Tactile, acoustic and vestibular systems sum to elicit the startle reflex. Neuroscience & Biobehavioral Reviews, 26, 1–11.

Yoder, R.M. & Taube, J.S. (2009) Head direction cell activity in mice: robust directional signal depends on intact otolith organs. J Neurosci, 29, 1061–1076.

Zugaro, M.B., Arleo, A., Berthoz, A. & Wiener, S.I. (2003) Rapid Spatial Reorientation and Head Direction Cells. The Journal of Neuroscience, 23, 3478–3482.

